# BRD4-mediated repression of p53 is a target for combination therapy in AML

**DOI:** 10.1101/2020.09.16.291930

**Authors:** Anne-Louise Latif, Ashley Newcombe, Sha Li, Kathryn Gilroy, Neil Robertson, Xue Lei, Helen Stewart, John Cole, Maria Terradas Terradas, Loveena Rishi, Lynn McGarry, Claire McKeeve, Claire Reid, William Clark, Joana Campos, Kristina Kirschner, Andrew Davis, Jonathan Lopez, Jun-Ichi Sakamaki, Jennifer Morton, Kevin M. Ryan, Stephen Tait, Sheela Abraham, Tessa Holyoake, Brian Higgins, Xu Huang, Karen Blyth, Mhairi Copland, Tim Chevassut, Karen Keeshan, Peter D. Adams

## Abstract

Acute Myeloid Leukemia (AML) is a typically-lethal molecularly heterogeneous disease, with few broad-spectrum therapeutic targets. Unusually, most AML retain wild-type *TP53*, encoding the pro-apoptotic tumor suppressor p53. MDM2 inhibitors (MDM2i), which activate wild-type p53, and BET inhibitors (BETi), targeting the BET-family co-activator BRD4, both show encouraging pre-clinical activity, but limited clinical activity as single agents. Here, we report synergistic toxicity of combined MDM2i and BETi towards AML cell lines, primary human blasts and mouse models, resulting from BETi’s ability to evict an unexpected repressive form of BRD4 from p53 target genes, and hence potentiate MDM2i-induced p53 activation. These results indicate that wild-type *TP53* and a transcriptional repressor function of BRD4 together represent a potential broad-spectrum synthetic therapeutic vulnerability for AML.

## Introduction

Despite numerous advances in the knowledge of the molecular landscape of Acute Myeloid Leukemia (AML), there remains an unmet need to improve clinical outcomes; 5-year survival rates in 2019 for adults diagnosed with AML remain below 30% (Watts and Nimer, 2018). AML is a genetically and epigenetically heterogeneous disease, characterized by recurrent but diverse chromosomal structural changes and genetic mutations associated with functionally distinct sub-groups (Klco et al., 2013; Papaemmanuil et al., 2016). In addition to disease heterogeneity between patients, marked sub-clonal heterogeneity has also been observed within individual AML patients (de Boer et al., 2018). To date these disease features have posed a significant obstacle to finding novel targeted agents with a broad therapeutic reach.

One key unifying feature of AML, that could potentially be exploited to benefit many patients, is that the majority of cases exhibit wild-type *TP53* (Klco et al., 2013; Papaemmanuil et al., 2016). In AML with wild-type *TP53*, the p53 tumour suppressor protein is commonly held functionally inert through dysregulation of the ARF-MDM2/4 axis, culminating in inactivation of p53 by its negative regulators, MDM2 and MDM4 (Prokocimer et al., 2017). Drugs and small molecules have been developed that can activate p53 in cells expressing the wild-type gene (Khoo et al., 2014), with the goal of unleashing p53’s potent tumor suppressive functions. Clinical grade MDM2 inhibitors (MDM2i) have been tested in the clinic, for example RG7112 in hematological and solid tumors, with some encouraging, but limited, responses (Andreeff et al., 2016; Khoo et al., 2014). Nevertheless, in pre-clinical studies, MDM2i cooperated with “standard-of-care” therapies, daunorubicin and cytarabine, to eradicate AML (Maganti et al., 2018); and the BCL2 inhibitor (Venetoclax) and MDM2i (Idasanutlin/RG7388 [a clinical grade RG7112 derivative with improved potency, selectivity and bioavailability] (Ding et al., 2013)) are also synthetic lethal in AML (Pan et al., 2017). Other combination strategies have also demonstrated the utility of targeting wild-type p53 in AML (Minzel et al., 2018). Consistent with these studies, Andreeff *et al* have proposed that use of MDM2i to activate p53 will likely realize more benefit in combination therapies (Andreeff et al., 2016).

Bromodomain-containing protein 4 (BRD4) is a member of the bromodomain and extraterminal (BET) family proteins, characterized by two N-terminal bromodomains and an extraterminal domain (Roe and Vakoc, 2016). BRD4 has been shown to play a role in the activation of genes involved in cell growth - most notably *c-MYC* - through binding to acetylated histones and transcription factors, to which BRD4 then recruits transcriptional regulators, such as positive transcription elongation factor b (P-TEFb) and Mediator complex (Roe and Vakoc, 2016). Although *c-MYC* translocations or mutations are not common in AML, the activation of *c-MYC* by multiple up-stream leukemic genetic aberrations has been recognized as a key hub in driving leukemogenesis (Delgado and Leon, 2010). Pre-clinical data has demonstrated that inhibition of BRD4 has efficacy across a range of AML subtypes (Dawson et al., 2011; Filippakopoulos et al., 2010; Zuber et al., 2011). Indeed, BET inhibitors (BETi) have entered early phase clinical trials for AML. However, despite promising pre-clinical activity, their efficacy in treating AML as single agents has been modest (Amorim et al., 2016; Berthon et al., 2016; Chaidos et al., 2015; Dombret and Gardin, 2016), and as such it is likely that, like MDM2i, their strength lies in rational combination therapies.

In sum, both MDM2i and BETi have been considered as therapies for AML, but on their own have shown limited clinical activity (Amorim et al., 2016; Andreeff et al., 2016; Berthon et al., 2016; Chaidos et al., 2015; Dombret and Gardin, 2016; Khoo et al., 2014). Given that both drugs can, in principle, target a broad spectrum of AML molecular subtypes and the two drugs have distinct modes of action, we set out to test the hypothesis that the concomitant reactivation of p53 and inhibition of BET family proteins, using MDM2i and BETi, could synergise to kill AML cells. Here we present data showing superior efficacy of the drug combination over the single agents in genetically heterogenous AML cell lines, primary AML samples, and two relevant mouse models. We present mechanistic data demonstrating how this efficacious drug combination co-operates to induce pro-apoptotic p53 target genes.

## Results

### BETi enhance the killing of human AML cells by MDM2i in a p53-dependent manner

Initial experiments to assess potential synergy of the MDM2i and BETi combination were performed in a panel of primary AML cells from 15 heterogeneous AML patients. These patients had a median age of diagnosis of 60 years (range 31 to 78 years). Based on their non-complex karyotype we expect the majority to retain wild-type *TP53* (Klco et al., 2013), and this was confirmed for the 4 samples for which DNA sequencing data was available (Supplementary Figure 1A). In these initial *in vitro* studies, for BETi we used CPI203 (Constellation Pharmaceuticals), and for MDM2i we used nutlin-3 (Sigma). CPI203 is a potent BETi with an IC50 of 37nM for inhibition of BRD4 (Moros et al., 2014). Like other BETi, CPI203 represses expression of oncoproteins, such as c-MYC (Moros et al., 2014). CPI203 is a pre-clinical tool compound version of CPI0610, another Constellation BETi already being tested in human AML (Albrecht et al., 2016). Nutlin-3 is a potent inhibitor of the interaction between p53 and its negative regulator MDM2 (Vassilev et al., 2004), thereby activating p53 to inhibit cell proliferation and tumorigenesis in models harboring wild-type *TP53*. As an assessment of potential drug synergy, we measured cell viability reflected in ATP levels (CellTitre-Glo, Promega). Across all 15 patient samples, the combination of CPI203 and nutlin-3 was significantly more efficacious than either drug alone (Figure 1A). To more quantitatively assess drug interactions, we calculated the coefficient of drug interaction (CDI) for each sample; CDI <1, =1 or >1 indicates that the drugs are synergistic, additive or antagonistic, respectively. The majority of the samples showed a trend towards greater than additive killing by the combination (x axis < 1), compared to the single drugs (Figure 1B). For 4 of the samples, this synergistic interaction was determined to be significant (p<0.05). Due to limited sample availability, *TP53* mutation status could only be determined by DNA sequencing for 4 of the human patient samples. Of these, all retained wild-type *TP53* and 3 of them showed at least a trend towards synergy. These results in primary patient samples point to synergistic toxicity of the BETi and MDM2i combination against a substantial proportion of primary human AML.

**Figure 1.**
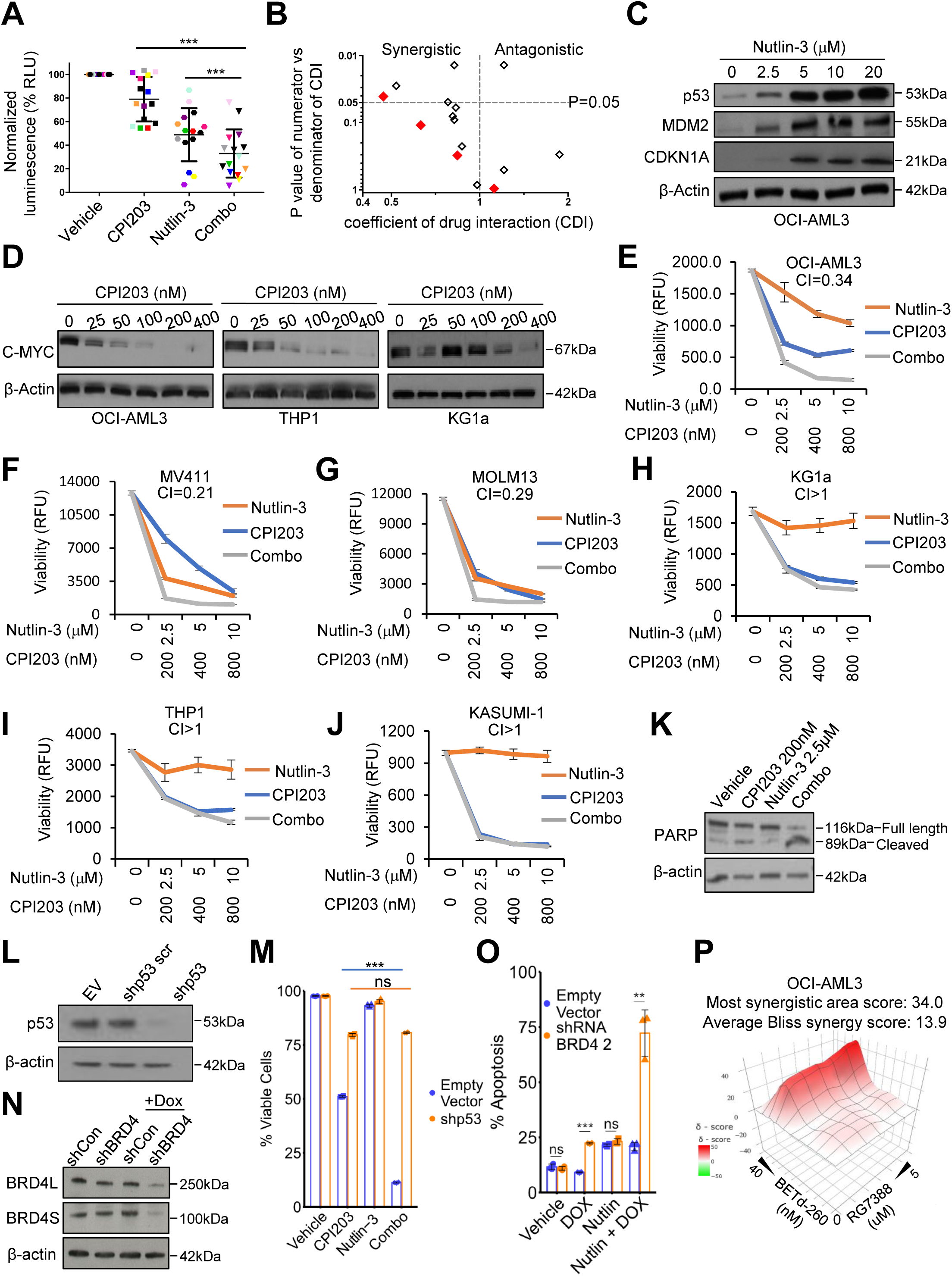
MDM2 and BET inhibitors are synergistically lethal to primary human AML blasts and AML cell lines with wild-type *TP53*. **A.** Primary human patient AML blasts (n=15) were treated with indicated drugs (2.5μM nutlin-3 alone, 200nM CPI203 alone, and the two in combination) and cell viability was assessed by CellTiter-Glo assay after 48h (***=p≤0.001, two-tailed unpaired t-test) **B.** Scatter plot of coefficient of drug interaction (CDI) and synergy significance in primary human patient blasts from A. CDI was calculated as follows: CDI=AB/(AxB). AB is the fraction of cells surviving (0-1.0) in the combination of drugs (nutlin-3+CPI203); A (nutlin-3) and B (CPI203) are the fraction of cells surviving in each of the single drugs. CDI value (x-axis) <1 (left), =1, or >1 (right) indicates that the drugs are synergistic, additive or antagonistic, respectively. p value (y-axis) ≤0.05 indicates significance (top). The p value was generated comparing AB and AxB (***=p≤0.001, two-tailed unpaired t-test). Red diamonds, *TP53* wild-type. Open diamonds, *TP53* status unknown. **C.** Western blots performed on the OCI-AML3 cell lines assessing expression of p53, MDM2, and CDKN1A after 24 hours of drug treatment with increasing doses of nutlin-3. **D.** Western blots performed on the OCI-AML3, THP1 and KG1a cell lines, assessing the expression of C-MYC after 24 hours in indicated doses of CPI203. **E.** OCI-AML3 cell viability (each treatment in triplicate) was assessed by resazurin assay after 72 hours, using a treatment ratio of CPI203:nutlin-3 of 1:12.5. Mean +/− Standard deviation of 3 independent replicates is shown. **F.** MV411 cell viability (each treatment in triplicate) was assessed by resazurin assay after 72hours, using a treatment ratio of CPI203:nutlin-3 of 1:12.5. Mean +/− Standard deviation of 3 independent replicates is shown. **G.** MOLM13 cell viability (each treatment in triplicate) was assessed by resazurin assay after 72hours, using a treatment ratio of CPI203:nutlin-3 of 1:12.5. Mean +/− Standard deviation of 3 independent replicates is shown. **H.** KG1a cell viability (each treatment in triplicate) was assessed by resazurin assay after 72 hours, using a treatment ratio of CPI203:nutlin-3 of 1:12.5. Mean +/− Standard deviation of 3 independent replicates is shown. **I.** THP1 cell viability (each treatment in triplicate) was assessed by resazurin assay after 72 hours, using a treatment ratio of CPI203:nutlin-3 of 1:12.5. Mean +/− Standard deviation of 3 independent replicates is shown. **J.** KASUMI-1 cell viability (each treatment in triplicate) was assessed by resazurin assay after 72 hours, using a treatment ratio of CPI203:nutlin-3 of 1:12.5. Mean +/− Standard deviation of 3 independent replicates is shown. **K.** Cleavage of PARP as assessed by Western blot analysis according to treatment condition after 24 hours treatment of the OCI-AML3 cell line. **L.** Western blot assessing expression of p53 in the OCI-AML3 cell line, harboring empty vector, scrambled shRNA p53 or shRNA p53. **M.** Cell viability as assessed by trypan blue in OCI-AML3 cells harboring empty vector or shRNA p53, treated for 72 hours with vehicle, 200nM CPI203, 2.5μM nutlin-3 or the drug combination (***=p≤0.001, two-tailed unpaired t-test, n=3, Means +/− SD are shown). **N.** Western blot analysis of shRNA-mediated knock down of BRD4 (shRNA induced by 0.5ug/ml doxycycline for 72 hours). **O.** FACS analysis of apoptosis (annexin V and PI) in OCI-AML3 cells expressing empty-vector or shRNA BRD4 (doxycycline-inducible), treated with vehicle, doxycycline, nutlin-3 and the combination of nutlin-3 and doxycycline for 72 hours (***=p≤0.001, **=p≤0.01, two-tailed unpaired t-test, n=3, Means +/− SD are shown). **P.** Excess Over Bliss plot showing synergistic effects between RG7388 and BETd-260 in OCI-AML3 cells. Cell viability (each treatment in quadruplicate) was assessed by CellTiter-Glo after 24 hours. Bliss synergy scores were indicated.

To extend these results obtained in primary AML blasts and to be able to perform more rigorous mechanistic analyses, we tested synergy between nutlin-3 and CPI203 in a panel of 3 human AML cell lines expressing wild-type p53 (OCI-AML3, MOLM13, MV411). To begin to assess the p53 dependence of any effect, we also tested 3 cell lines with mutant *TP53* (THP1, KG1a, Kasumi1). Irrespective of *TP53* status, these AML cell lines reflect diverse AML subtypes. For example, OCI-AML3 harbor *DNMT3A* and *NPM1* mutations, MOLM13 and MV411 both contain mutant *FLT3-ITD*, MOLM13 expresses an MLL-AF9 fusion oncoprotein, while MV411 expresses the MLL-AF4 fusion oncoprotein ((Matsuo et al., 1997) and ATCC). We confirmed initially that the drugs engaged their molecular targets, reflected in upregulation of p53 and CDKN1A (p21) by nutlin-3, and down regulation of c-MYC by CPI203 (irrespective of *TP53* status) (Figure 1C, D). To compare the effects of the single drugs and combination on viability of these 6 cell lines, we used resazurin (a metabolic viability sensor) across a range of fixed drug dose ratios, enabling drug interaction to be quantitatively assessed by the combination index (CI) method, where CI<1 is generally considered as synergy (Chou and Talalay, 1984). Consistent with previous studies in OCI-AML3 cells (Stewart et al., 2013), the drug combination demonstrated apparent synergy in wild-type *TP53* cell lines (Figure 1E-G and Supplementary Figure 1B-D). In the *TP53* wild-type OCI-AML3 cell line, CPI203 and nutlin-3 were designated synergistic from CI 0.34-0.52 at different drug ratios (Figure 1E and Supplementary Figure 1B-D). Similarly, MV411 and MOLM13 showed CI 0.21 and 0.29, respectively (Figure 1F, G). There was no benefit in using the combination treatment on the *TP53* mutated cell lines (Figure 1H-J) (CI for all mutant *TP53* lines > 1). To confirm these results we also calculated drug synergy by the Bliss method (Ianevski et al., 2020). This also showed substantially greater and more consistent synergy (Bliss score >0) of the combination in *TP53* wild-type cells, across a range of drug doses (Supplementary Figure 1E-J). The results of the cell viability studies in the *TP53* wild-type OCI-AML3 cell line were corroborated by annexin V/propidium iodide flow cytometry assay. We observed marked enhancement of apoptosis specifically (annexin V positive) and/or cell death by apoptosis or other mechanisms (annexin V/PI positive), compared to either single drug alone (Supplementary Figure 1K-M). Enhanced killing of the combination-treated cells was accompanied by a clear increase in cleaved PARP (Figure 1K), a marker of pro-apoptotic caspase activation, showing that the drug combination induces apoptosis in these *TP53* wild-type AML cells.

Confirming a requirement for wild-type *TP53* for the drug combination to induce super-additive cell death, we knocked down p53 using shRNA in the *TP53* wild-type OCI-AML3 cell line which abrogated the benefit of the drug combination (Figure 1L, M). To similarly confirm that the effects of CPI203 were mediated via inhibition of a BET family member, we asked whether the enhanced killing by the MDM2i/BETi combination was recapitulated by MDM2i together with BRD4 knock down. Indeed, inducible shRNA-mediated knock down of BRD4 markedly potentiated killing by MDM2i (Figure 1N, O). In addition, we used BET PROTACs to selectively and completely induce degradation of BRD4 in cells (Supplementary Figure 1N)(Lu et al., 2015; Zhou et al., 2018). Consistent with BETi, BET degraders (BETd-260 and ARV-825) synergize with MDM2i to suppress cell viability in 3 wild-type *TP53* cell lines (OCI-AML3, MOLM13, MV411) (Figure 1P and Supplementary Figure 1O-S). Taken together, in primary human AML blasts and human AML cell lines, we conclude that, in AML expressing wild-type p53, BETi inhibit BRD4 to enhance cell killing by MDM2i.

### BETi and MDM2i cooperate to eradicate AML in mouse models

Next, we asked whether the drug combination shows superior anti-leukemic activity to the single agents, in two murine models of AML. *Trib2* is an oncogene capable of causing AML in mice through down regulation of the transcription factor C/EBPα, a gene that is mutated in 15% of cases of AML in humans (Keeshan et al., 2006). Following confirmation by PCR and Sanger sequencing that blasts from mice with AML driven by Trib2 expressed wild-type p53 (data not shown), we first confirmed superior eradication of the murine leukemic cells by the drug combination compared to the two single drugs *in vitro* (Figure 2A-C), recapitulating the effect in human cells. For the purposes of subsequent *in vivo* work wanted to use clinical grade drugs. We used the clinical grade BETi CPI0610 (Constellation Pharmaceuticals) and MDM2i RG7112 (Roche Pharmaceuticals). RG7112 has been tested as a single agent in relapsed refractory AML (Andreeff et al., 2016), showing modest activity, and is better suited to studies with murine AML than human-optimized RG7388 (Ding et al., 2013). CPI0610 is currently being tested in early stage human clinical trials and has been reported to be well-tolerated (Bankar and Gupta, 2020). For our experiments, we also identified well-tolerated drug doses in mice, based on a 21-day drug pilot experiment, assessing weight loss and myeloid cell counts (in bone marrow), B cells (in spleen) and T cells (in thymus) in non-leukemic normal healthy mice (Supplementary Figure 2A, B). Forty C57BL/6 mice (ten mice each for vehicle, both single drugs and the combination) were sub-lethally irradiated and 0.85×10^6^ Trib2 AML blasts injected via their tail veins. Myeloblasts expressed GFP from the same retroviral construct as Trib2 for disease tracking. 21 days of drug treatment was commenced in all leukemic mice, post confirmation of comparable disease engraftment between groups (data not shown). Three mice from each treatment group were sacrificed after the first 48hrs of drug treatment, GFP+ blasts recovered from bone marrow, RNA extracted and qPCR performed to demonstrate that RG7112 increased expression of *CDKN1A* and CPI0610 reduced levels of *c-MYC* (Figure 2D-E), as expected if the drugs engage their respective targets. After 21 days of treatment, all remaining mice were sacrificed (n= 7 for each group, except n=6 for vehicle because 1 succumbed to disease 15 days post engraftment) and abundance of GFP+ AML blasts in bone marrow determined by FACS. The drug combination demonstrated significantly enhanced anti-leukemic activity compared to either single drug alone, measured in bone marrow (Figure 2F-G), spleen, and thymus (Supplementary Figure 2C). In bone marrow and spleen the CDI was <1 (0.017 and 0.38 respectively), indicating marked synergy *in vivo*.

**Figure 2.**
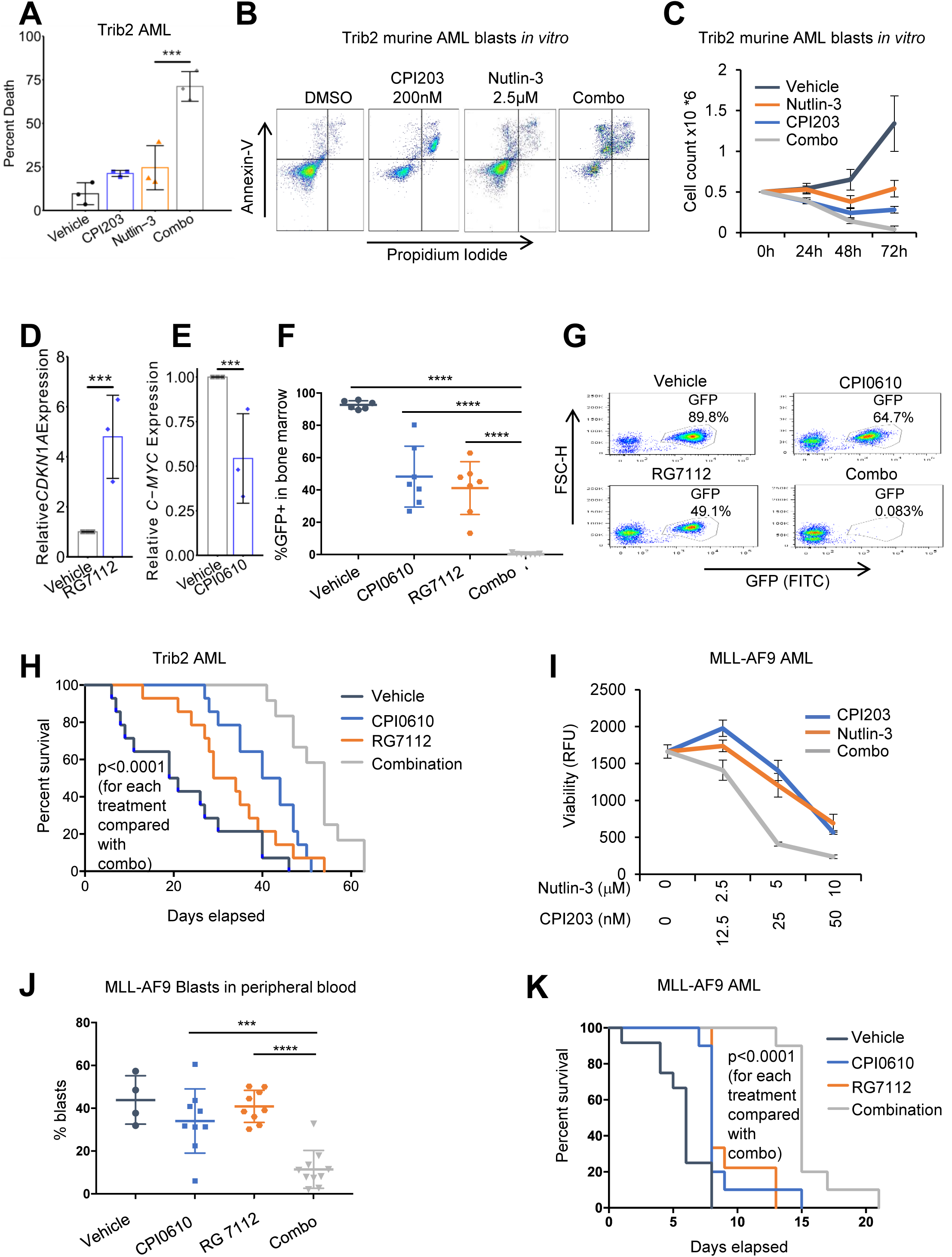
MDM2 and BET inhibitors cooperate to eradicate AML in *in vivo* mouse models. **A.** Primary murine Trib2 AML blasts, treated *in vitro* for 72 hours with vehicle, 200nM CPI203, 2.5μM nutlin-3 or the drug combination. Cell death assessed by flow cytometry using Annexin-V and PI staining (***=p≤0.001, two-tailed unpaired t-test, n=3, Means +/− SD are shown). **B.** A representative example of the data summarized in Figure 2A (Gating strategy for sorting GFP Positive cells was live cells>cell profile>single cells>GFP positive cells.). **C.** A time course of viable cell counts as assessed by trypan blue staining, of primary murine Trib2 AML blasts, according to treatment condition (Error bars are SEM, n=3, Means +/− SD are shown). **D.** qPCR assessment of CDKN1A expression in GFP+ blasts recovered from bone marrow from three mice sacrificed after 48hrs treatment with RG7112 relative to vehicle (**=p≤0.01, two-tailed unpaired t-test, n=3, Means +/− SD are shown). **E.** qPCR assessment of C-MYC expression in GFP+ blasts recovered from bone marrow from three mice sacrificed after 48hrs treatment with CPI0610 relative to vehicle (***=p≤0.001, two-tailed unpaired t-test, n=3, Means +/− SD are shown). **F.** Assessment of the disease burden in the bone marrow according to treatment condition, at the end of 21 days of treatment *in vivo*; disease burden assessed by flow cytometry to measure percent GFP+ blasts (****=p≤0.0001, two-tailed unpaired t-test, from left to right: n=6, 7, 7, 7, Means +/− SD are shown). **G.** A representative example of the data summarized in Figure 2F (Gating strategy for sorting GFP Positive cells was live cells>cell profile>single cells>GFP positive cells.). **H.** A survival analysis of mice transplanted with Trib2-driven AML according to treatment condition (****=p≤0.0001, from Kaplan-Meier estimator). **I.** Primary murine MLL-AF9 AML blasts treated *in vitro* with vehicle, 200nM CPI203, 2.5μM nutlin-3 and the drug combination for 72 hours. Cell viability assessed by resazurin. (Means +/− SD are shown, n=3) **J.** Analysis of disease burden in peripheral blood, based on percent CD11b^low^ Gr1+ blasts, after 7 days of treatment, according to drug condition (***=p≤0.001,****=p≤0.0001, two-tailed t-test, from left to right, n=4, 9, 9, 10, Means +/− SD are shown). **K.** A survival analysis of mice transplanted with MLL-AF9-driven AML, according to treatment condition (****=p≤0.0001, from Kaplan-Meier estimator).

Following demonstration that the drug combination was tolerable and efficacious at the end of the treatment period, we assessed if the drug combination could confer a survival advantage over the single agents in mice with Trib2-driven AML. Sub-lethally irradiated cohorts of mice were again injected with GFP+ Trib2 AML blasts and, following comparable disease engraftment between groups (Supplementary Figure 2D), were treated with vehicle, single drugs or combination for 21 days. FACS analysis of peripheral blood at the end of drug treatment confirmed the strong synergy of the drug combination in eradication of GFP+ blasts (CDI=0.031, Supplementary Figure 2E). After completion of the 21-day drug cycle, mice were closely monitored for signs of leukemia and culled at an ethical endpoint. Although the combination-treated mice did ultimately succumb to disease after cessation of treatment, indicative of low level residual disease, the combination-treated mice did have a statistically significant survival advantage compared with the single agent-treated mice (Figure 2H). This underscores the anti-leukemic activity of the combination in this model.

To confirm that these findings were not confined to this leukemia, given the heterogeneous nature of AML, we sought to demonstrate that the drug combination could also elicit improved killing in a distinct AML mouse model, namely the MLL-AF9 mouse model. This aggressive AML, driven by a *KMT2A* translocation occurring in ∼5% of human adult AML, is well-established and frequently utilized for testing novel drugs for AML (Somervaille and Cleary, 2006). We first confirmed by DNA sequencing that murine MLL-AF9 cells harbored wild-type p53 (data not shown), and that at most drug ratios the drug combination enhanced killing of MLL-AF9 myeloblasts *in vitro* (Figure 2I). Sub-lethally irradiated cohorts of mice were injected with GFP+ MLL-AF9 AML blasts and cohorts of comparable disease burden (Supplementary Figure 2F) were treated with vehicle, single drugs or combination for 21 days. Analysis of disease burden in peripheral blood after 7 days of treatment, based on GFP+ blasts and CD11b^low^ Gr1-expressing immature blasts known to expand in AML (Somervaille and Cleary, 2006), demonstrated the superiority of the drug combination over single agents in disease suppression (CDI=0.36, Figure 2J and Supplementary Figure 2G). Moreover, although at this high dose of leukemic cells (200,000 per mouse) the vehicle-treated mice succumbed to disease in less than 2 weeks, the drug combination-treated mice showed a robust and highly statistically significant survival advantage compared with the single agent-treated mice in the MLL-AF9 model (Figure 2K). As expected, the 4 treatment cohorts showed comparable disease burden at the time of cull based on similar disease symptoms (Supplementary Figure 2H). We conclude that in two different p53 wild-type mouse models of AML, the drug combination is superior to the single agents in suppressing disease and extending survival.

### BETi potentiate activation of p53 by MDM2i

We set out to define the basis of synergy between BETi and MDM2i in killing AML cells. In the OCI-AML3 cell line, treatment with single agent CPI203 reduced the protein abundance of c-MYC as expected, but this was not substantially further reduced by the drug combination (Figure 3A). Likewise, nutlin-3 modestly increased abundance of p53, but again this was not enhanced by the drug combination (Figure 3A). Therefore, synergy does not appear to result from a concerted effect of the drugs on stabilization of p53 nor repression of c-MYC. Previously, a combination of MDM2i and the BCL2 inhibitor Navitoclax has been shown to synergize in killing AML cells (Pan et al., 2017). Since BETi downregulate expression of BCL2 and upregulate expression of BCL2 inhibitor BIM (Dawson et al., 2011; Xu et al., 2016), we wondered whether BETi-mediated downregulation of BCL2 activity contributes to synergy between BETi and MDM2i. We confirmed that BETi CPI203 downregulates BCL2 (Figure 3B). However, ectopic expression of BCL2 did not suppress killing by the combination (Figure 3C, D). These results do not eliminate a role for regulation of BCL2 family proteins (including BIM) in drug combination-induced AML cell killing, but do indicate that BETi-mediated repression of BCL2 is not necessary for killing by the combination.

**Figure 3.**
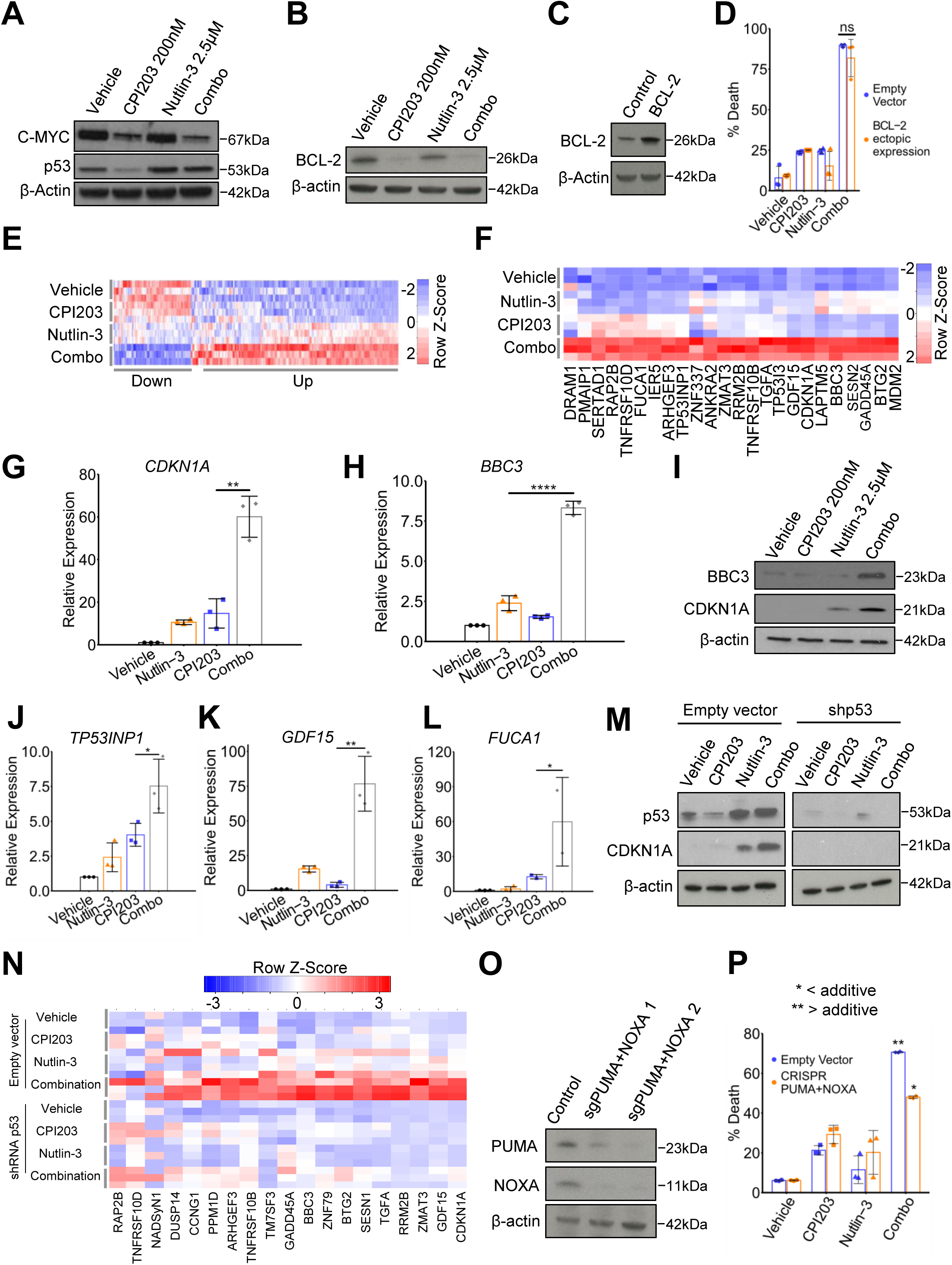
BET inhibitors potentiate activation of p53 target genes by p53. **A.** Western blot of C-MYC and p53 in OCI-AML3 cells treated for 24 hours with vehicle, 200nM CPI203, 2.5μM nutlin-3 and the drug combination. **B.** Western blot for BCL-2 in OCI-AML3 cells treated for 24 hours with vehicle, 200nM CPI203, 2.5μM nutlin-3 and the drug combination. **C.** Western blot for BCL-2 comparing the control OCI-AML3 and cells over-expressing BCL-2. **D.** Cell killing as assessed by flow cytometry using Annexin-V and PI staining of OCI-AML3 control (empty vector) cells versus OCI-AML3 over-expressing BCL-2, according to treatment condition (p-value >0.05 by two tailed unpaired t-test, Means +/− SD are shown, n=3). **E.** A heat map of genes synergistically regulated by the drug combination determined by RNA-seq of OCI-AML3 cells according to treatment condition (24hrs). Genes synergistically up-regulated (Up) by the drug combination were rigorously identified as those where C/(A+B) => 1.25 and combination FPKM > DMSO FPKM, and synergistically down-regulated genes (Down) where C/(A+B) => 1.25 and combination FPKM < DMSO FPKM (where C= combination FPKM – DMSO FPKM; A= nutlin-3 FPKM – DMSO FPKM; B= CPI203 FPKM – DMSO FPKM). **F.** A heat map of 24 high confidence p53 target genes that are synergistically up-regulated or down-regulated in OCI-AML3 according to treatment condition. **G.** qPCR analysis of *CDKN1A* expression in OCI-AML3 under indicated treatments for 24h (**=p≤0.01, two tailed unpaired t-test, Means +/− SD are shown, n=3). **H.** qPCR analysis of *BBC3* expression in OCI-AML3 under indicated treatments for 24h (****=p≤0.0001, two tailed unpaired t-test, Means +/− SD are shown, n=3). **I.** Western blot analysis of p53 targets BBC3 and CDKN1A in OCI-AML3 according to treatment condition. **J.** qPCR expression analysis of *TP53INP1* in OCI-AML3 under indicated treatments for 24h (*=p≤0.05, two tailed unpaired t-test, Means +/− SD are shown, n=3). **K.** qPCR expression analysis of *GDF15* in OCI-AML3 under indicated treatments for 24h (**=p≤0.01, two tailed unpaired t-test, Means +/− SD are shown, n=3). **L.** qPCR expression analysis of *FUCA1* in OCI-AML3 under indicated treatments for 24h (*=p≤0.05, two tailed unpaired t-test, Means +/− SD are shown, n=3). **M.** Western blot analysis of p53 and CDKN1A in OCI-AML3 cells and OCI-AML3 cells harboring shRNA p53 according to treatment condition. **N.** A heat map of an RNA-seq data of 19 high confidence p53 target genes that are synergistically up-regulated by the drug combination in control (empty-vector) OCI-AML3 cells, versus shRNA p53 OCI-AML3 cells, according to treatment condition. **O.** Western blot analysis of BBC3 and NOXA in control (empty vector) OCI-AML3 cells, and OCI-AML3 cells harboring CAS9 and sgRNAs against PUMA and NOXA. **P.** FACS analysis of cell death (annexin VI and PI) in control OCI-AML3 cells or cells in which PUMA and NOXA were knocked out by CRISPR/CAS9, under indicated conditions for 72 hours (*=<additive. **=>additive, two tailed unpaired t-test, Means +/− SD are shown, n=3).

To obtain unbiased insight into the potential basis of this drug synergy, we set out to compare the gene expression profiles of OCI-AML3 cells treated with either vehicle, CPI203 (200nM), nutlin-3 (2.5μM) or the drug combination. OCI-AML3 cells were treated with vehicle, CPI203 200nM, nutlin-3 2.5μM and the drug combination for 24 hours, then RNA was extracted and analyzed by RNA-seq. Principal Component Analysis confirmed the expected difference between the drug treatments at the gene expression level (Supplementary Figure 3A). Following treatment with CPI203, either as a single agent or as part of the drug combination, ∼6,100 coding genes significantly changed expression relative to vehicle (Supplementary Figure 3B-D). A much smaller number of genes (174) significantly changed expression following treatment with nutlin-3 only (Supplementary Figure 3B-D). This was anticipated given the broad effects of BET inhibition, at least in part a consequence of down-regulation of c-MYC, compared with a more restricted number of known p53 target genes (Dawson et al., 2011; Filippakopoulos et al., 2010; Zuber et al., 2011). We reasoned that synergistic killing by the drug combination might be underpinned by synergistic changes in gene expression. Indeed, visual analysis of a heatmap of significantly changed genes revealed clusters of apparently synergistically up and down-regulated genes (Supplementary Figure 3E). Quantitative analysis yielded 252 genes that were synergistically up-regulated and 94 genes that were synergistically down-regulated by the drug combination (Figure 3E and Supplementary Tables 1 and 2). Ingenuity Pathway Analysis (IPA) analysis showed that the synergistically down-regulated genes were enriched in cell cycle genes (Supplementary Figure 3F). The synergistically up-regulated genes were most significantly enriched in genes involved in the p53 pathway (Supplementary Figure 3F). Of 116 known high-confidence p53 target genes (Fischer, 2017), 24 were synergistically upregulated by the drug combination (Figure 3F and Supplementary Figure 3G). Synergistically up-regulated *TP53* target genes included *ANKRA2, ARHGEF3, BBC3 (PUMA), BTG2, CDKN1A, DRAM1, FUCA1, GADD45A, GDF15, IER5, LAPTM5, MDM2, PMAIP1, RAP2B, RRM2B, SERTAD1, SESN2, TGFA, TNFRSF10B, TNFRSF10D, TP53I3, TP53INP1, ZMAT3* and *ZNF337* (Figure 3F). Representative sequence tracks of p53 target genes *GDF15, CDKN1A* and *BBC3* are shown (Supplementary Figure 3H). Synergistic upregulation of p53 targets, *CDKN1A* and *BBC3*, by the drug combination was confirmed at the RNA level by qPCR and the protein level by western blot (Figure 3G-I), and at the RNA level for *GDF15, TP53INP1* and *FUCA1* (Figure 3J-L). We conclude that synergistic killing by the drug combination is associated with enhanced expression of p53 target genes.

To confirm that expression of these p53 target genes is indeed dependent on p53, we generated cells expressing shRNA to knock down p53 (or empty vector control cells) and treated with single or combination drugs (Figure 3M). p53-deficient cells failed to upregulate CDKN1A in response to nutlin-3 or combination (Figure 3M and Supplementary Figure 3I). Principal Component Analysis (PCA) of RNA-seq data confirmed that p53 knock down blunted the effect of the drug combination on the whole transcriptome (data not shown). Similar to the previous experiment, 214 genes were calculated to be synergistically upregulated by the drug combination in control cells. Of these, 204 were dependent on p53 for their synergistic upregulation (Supplementary Table 3). Of the 116 *bona fide* p53 target genes (Fischer, 2017), 19 were upregulated in control AML3 cells expressing p53 (12 of which were in common with the previous experiment (Figure 3F, 3N and Supplementary Figure 3J). None of these 19 p53 target genes was synergistically upregulated in drug combination-treated p53-deficient cells (Figure 3N). To dissect which p53 target genes are required for drug synergy in OCI-AML3 cells, we used CRISPR/Cas9 to generate derivatives of OCI-AML3 cells lacking *CDKN1A, BBC3* and *NOXA.* Consistent with the observation that drug combination synergy is linked to cell death (Figure 1), inactivation of CDKN1A, a well-known effector of p53-mediated cell cycle arrest but not apoptosis, did not affect synergy of the combination (Supplementary Figure 3K). More surprisingly, inactivation of pro-apoptotic p53-target genes *BBC3* and *NOXA* on their own did not significantly affect cell killing (data not shown). However, combined inactivation of *BBC3* and *NOXA* markedly blunted drug combination-induced cell killing (Figure 3O, P). Together, these results unexpectedly show that BET inhibition potentiates activation of p53. WT p53 is required for enhanced expression of p53 target genes by BET inhibition and at least two of these genes, *BBC3* and *NOXA*, are required for efficient synergistic killing of AML by combined BET and MDM2 inhibition.

### BET inhibition relieves BRD4-mediated repression of p53 target genes

We set out to define the molecular mechanism by which BET inhibition potentiates activation of p53. Induction of CDKN1A and BBC3 by nutlin-3 was also potentiated by BRD4 knock down, confirming that BETi acts, at least in part, by inhibition of BRD4 (Figure 4A, B). We initially considered three non-mutually exclusive testable possibilities for how BETi/inhibition of BRD4 potentiates activation of p53 by MDM2i. First, we postulated that the BETi might stabilize p53 target mRNAs. Second, we considered the possibility that BETi might, directly or indirectly, promote expression of known p53 activators that synergize with p53 stabilized by MDM2i to potentiate activation of the p53 pathway. Third, we hypothesized that BETi might promote p53 binding to its cognate target genes. To test whether BETi stabilized expression of p53 target mRNAs, OCI-AML3 cells were treated with vehicle, single drugs or combination and then 2 hours later with actinomycin D to inhibit new transcription. Abundance of p53 target mRNAs was determined by qPCR over a timecourse, allowing relative assessment of mRNA half-life. Figure 4C, D demonstrates that, compared to either single agent, the drug combination did not promote stability of *CDKN1A* or *BBC3* mRNAs.

**Figure 4.**
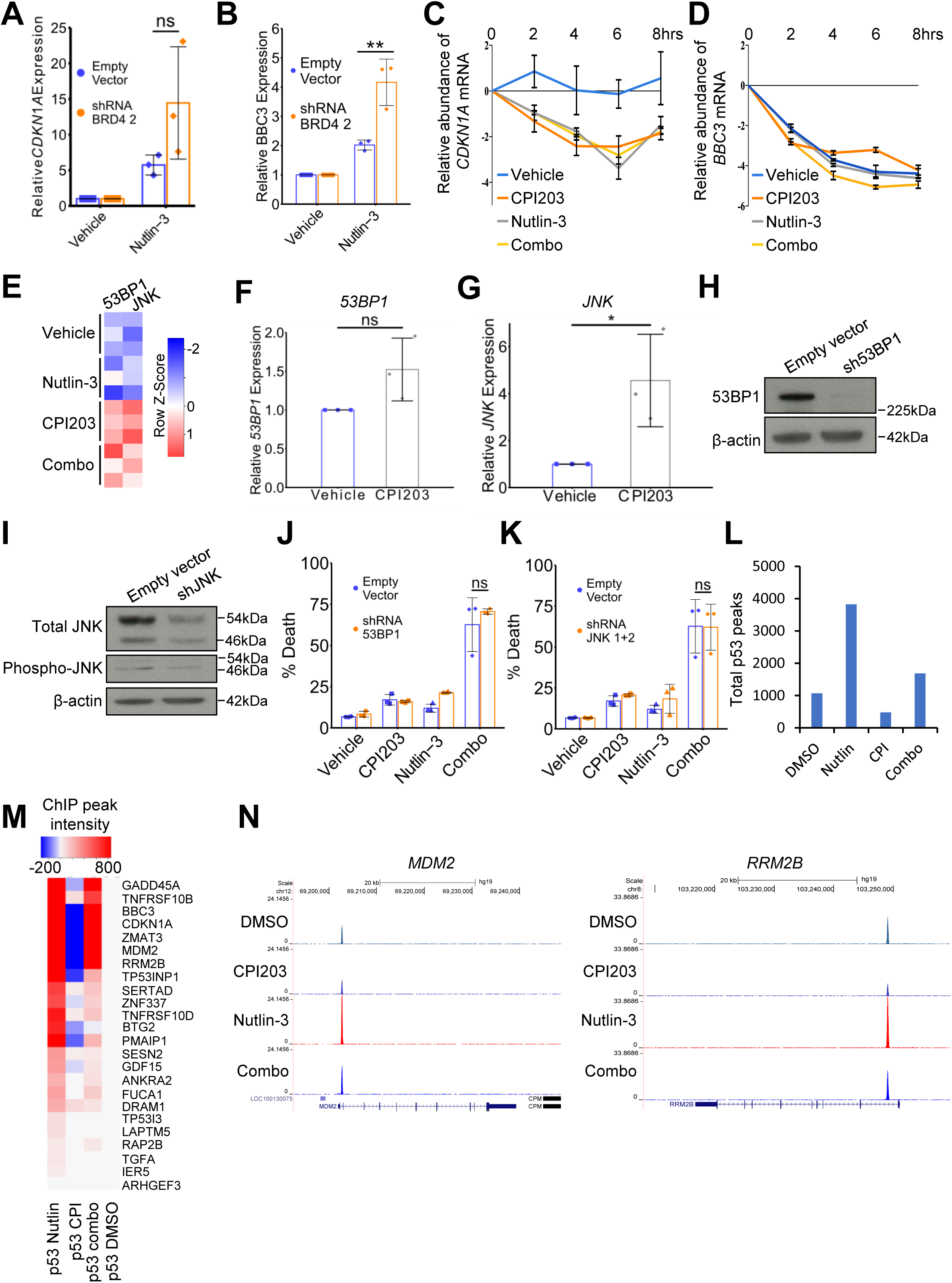
BET inhibitors do not stabilize p53 target mRNAs nor increase binding of p53 to target genes. **A.** qPCR assessment of expression of *CDKN1A* in control (empty-vector) OCI-AML3 cells versus OCI-AML3 cells harboring shRNA against *BRD4*, comparing treatment with vehicle and 2.5μM nutlin-3 (p-value >0.05 by two tailed unpaired t-test, Means +/− SD are shown, n=3). **B.** qPCR assessment of expression of *BBC3* in control (empty-vector) OCI-AML3 cells versus OCI-AML3 cells harboring shRNA against *BRD4*, comparing treatment with vehicle and 2.5μM nutlin-3 (two tailed unpaired t-test, Means +/− SD are shown, n=3, ** means p-value < 0.01). **C.** qPCR analysis of expression of *CDKN1A* in OCI-AML3 cells treated with Actinomycin D over the indicated time-course (0, 2, 4, 6, 8 hrs) and under the indicated drug treatments (Means +/− SD are shown, n=3). **D.** qPCR analysis of *BBC3* in OCI-AML3s treated with Actinomycin D over the indicated time-course (0, 2, 4, 6, 8 hrs) and under the indicated drug treatments (Means +/− SD are shown, n=3). **E.** Heat map showing expression of *JNK* and *53BP1* in OCI-AML3 cells after the indicated drug treatments for 24 hours. **F.** qPCR expression analysis of *53BP1* in OCI-AML3 cells after indicated drug treatments for 24h (two tailed unpaired t-test, Means +/− SD are shown, n=3). **G.** qPCR expression analysis of *JNK* in OCI-AML3 cells after indicated drug treatments for 24h (*=p≤0.05, two tailed unpaired t-test, Means +/− SD are shown, n=3). **H.** Western blot of 53BP1 in OCI-AML3 expressing either empty vector or shRNA *53BP1* **I.** Western blot of *JNK* in OCI-AML3 cells expressing either empty vector or shRNA *JNK* isoform 1 + isoform 2. **J.** FACS analysis of cell death (annexin VI and PI) in OCI-AML3 cells expressing empty-vector or shRNA *53BP1* after indicated drug treatments for 72 hours **K.** FACS analysis of apoptosis (annexin VI and PI) in OCI-AML3 cells expressing empty-vector or shRNA *JNK* isoform 1 + isoform 2 after indicated drug treatments for 72 hours **L.** Total number of p53 binding sites across the genome (ChIP-seq) after indicated drug treatments for 6 hours, present in at least 2 out of 3 replicates. **M.** Heat map of p53 binding at indicated genes in OCI-AML3 cells after indicated drug treatments for 6 hours. **N.** Sequence tracks of p53 binding at *MDM2* gene after indicated drug treatments for 6 hours. **O.** Sequence tracks of p53 binding at *RRM2B* gene after indicated drug treatments for 6 hours.

To test the role of selected p53 activators, we mined our RNA-seq data for known p53 activators upregulated by BETi. Based on analysis of RNA-seq data, at least two known p53 activators, *JNK* and *53BP1* (Cuella-Martin et al., 2016; Fuchs et al., 1998), were upregulated by BETi alone and the drug combination (Figure 4E). qPCR confirmed significant upregulation of *JNK* by BETi and an upward trend for *53BP1* (Figure 4F, G). To test a role for JNK or 53BP1 in activation of p53, we knocked them down by lentivirus-delivered shRNA (Figure 4H, I). However, neither was required for synergy of MDM2i and BETi (Figure 4J, K).

To test whether the drug combination promoted binding of p53 to its target genes compared to either single agent alone, we performed ChIP-seq to assess p53 binding across the whole genome in OCI-AML3 cells. Three independent replicates of OCI-AML3 cells were treated with vehicle control, CPI203 200nM, nutlin-3 2.5μM or the drug combination. Cells were harvested for ChIP 6hrs after drug treatment, since qPCR and western blot analysis showed that upregulation of p53 target genes was already detectable at this time (data not shown). Analysis of ChIP-seq data showed that the mean p53 peak width under all treatment conditions was ∼400bp (Supplementary Figure 4A). The overall genome distribution of sites under all conditions was similar and very few sites were unique to drug combination-treated cells (Supplementary Figure 4B, C). In fact, the drug combination tended to decrease the total number of p53 binding sites, compared to nutlin-3 alone (Figure 4L, Supplementary Figure 4C, D). Most of the 252 synergy up genes did not bind p53 in either nutlin-3 or combination-treated cells (Supplementary Figure 4E), suggesting that they are not direct p53 targets and their activation by nutlin-3 is indirect. For the 24 synergy up genes that are also *bona fide* p53 target genes, most bound p53 in the nutlin-3-treated cells, as expected, but this was not increased in combination-treated cells (Figure 4M). Instead, at p53 target genes, p53 binding tended to decrease in combination-treated cells compared to nutlin-3-treated cells (Figure 4M), in line with the global analysis (Figure 4L and Supplementary Figure 4C, D). Examination of the sequence tracks for individual genes, such as *MDM2* and *RRM2B*, confirmed the observation that BETi, in combination with nutlin-3, did not increase the binding of p53 to its target genes (Figure 4N, O). We ultimately conclude that BET inhibition does not activate p53 by stabilizing its target gene mRNAs, nor via promotion of activity of selected tested known p53 activators, nor by increasing p53’s binding to its target genes.

At this point, closer inspection of the RNA-seq data revealed that p53 was ranked as one of the top upstream regulators of genes differentially expressed by CPI203 (Supplementary Figure 5A). Focused analysis of p53 target genes directly showed that CPI203 alone was sufficient to increase expression of many p53 target genes in OCI-AML3 cells (Supplementary Figure 5B, C). For several p53 target genes this was confirmed by qPCR after treatment with CPI203 or knock down of BRD4 by lentivirus-encoded shRNA (Supplementary Figure 5D, E). Inhibition of BET family proteins with a potent PROTAC BET degrader BETd-260 strongly upregulated expression of p53 target gene p21 at the protein level (Supplementary Figures 1N and 5F) (Zhou et al., 2018). Since previous results established a critical role for BRD4 (Figure 1N, O and Figure 4A, B), these data raised the possibility that BRD4 might be a repressor of p53 target gene expression. To test this, we better characterized the effect of BETi and nutlin-3 on BRD4’s chromatin binding. We performed ChIP-seq analysis to determine genomic localization of BRD4 in control, single drug and combination-treated cells. In control OCI-AML3 cells, BRD4 reproducibly bound ∼9800 sites occupying ∼27Mb across the genome (Figure 5A, B and Supplementary Figure 6A). In nutlin-3-treated cells, a similar number of BRD4 binding sites and total genome occupancy was observed (Figure 5A, B and Supplementary Figure 6B), and the majority of binding sites overlapped between control and nutlin-3-treated cells (Supplementary Figure 6C). As expected, in BETi and combination-treated cells, the number of reproducible BRD4 binding sites was greatly decreased (Figure 5A, B and Supplementary Figure 6C-E). In both control and nutlin-3-treated cells the majority of the BRD4 binding sites were at annotated gene promoters (Figure 5C). Across all genes, in vehicle-treated cells BRD4 binding correlated with gene expression (Supplementary Figure 6F) and globally a loss of BRD4 on treatment with BETi was associated with a small but significant decrease in gene expression (Supplementary Figure 6G), consistent with the role of BRD4 as a global transcriptional activator (Roe and Vakoc, 2016).

**Figure 5.**
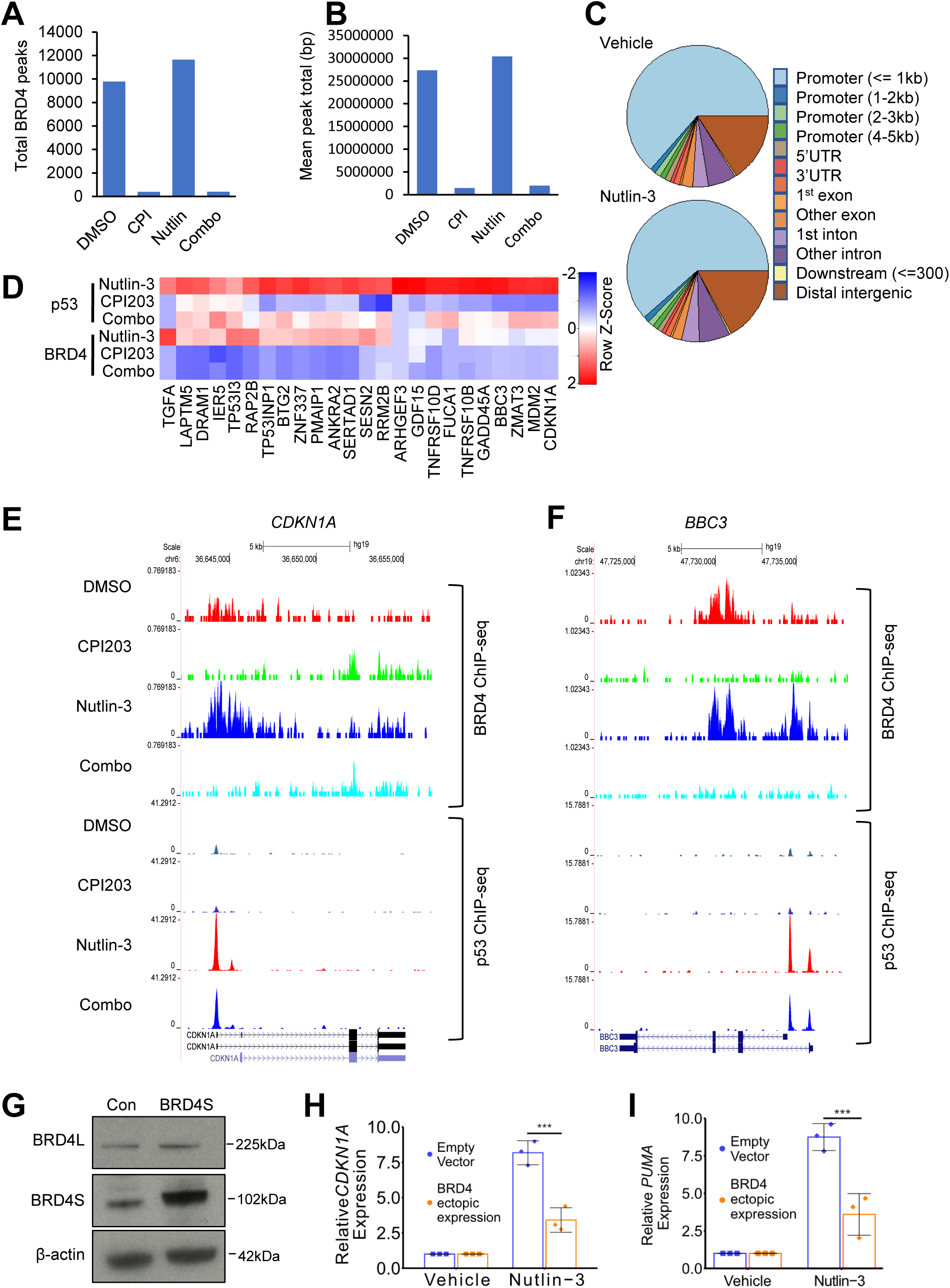
BRD4 represses p53 target genes. **A.** Total BRD4 peaks determined by ChIP-seq (present in both replicates) in OCI-AML3 cells after indicated drug treatments (6hrs). **B.** Mean total base pairs covered by BRD4 ChIP-seq peaks (peaks present in 2 out of 2 replicates) after the indicated drug treatments of OCI-AML3 cells. **C.** Genomic distribution of BRD4 binding sites identified by ChIP-seq (peaks present in 2 out of 2 replicates), after the indicated drug treatments of OCI-AML3 cells. **D.** Heat map showing BRD4 ChIP-seq (peaks present in 2 out of 2 replicates) at 24 p53 target genes, after the indicated drug treatments of OCI-AML3 cells. **E.** Sequence tracks of representative BRD4 and p53 binding at *CDKN1A,* after the indicated drug treatments of OCI-AML3 cells. **F.** Sequence tracks of representative BRD4 and p53 binding at *BBC3,* after the indicated drug treatments of OCI-AML3 cells. **G.** Western blot for BRD4 in OCI-AML3 cells ectopically expressing the short isoform of BRD4 (BRD4S). BRD4L is the long isoform. **H.** qPCR analysis of *CDKN1A* in OCI-AML3 cells ectopically expressing BRD4S, in absence or presence of nutlin-3 (***=p≤0.001, two tailed unpaired t-test, Means +/− SD are shown, n=3). **I.** qPCR analysis of *BBC3A* in OCI-AML3 cells ectopically expressing BRD4S, in absence or presence of nutlin-3 (***=p≤0.001, two tailed unpaired t-test, Means +/− SD are shown, n=3).

To evaluate the role of BRD4 in potentiation of p53 target gene expression by BETi, we considered BRD4 binding in nutlin-3 vs combination-treated cells. For the 24 synergy up genes that are also *bona fide* p53 target genes (Figure 4F), 22 bound BRD4 in nutlin-3-treated cells but only 2 bound BRD4 in combination-treated cells (Figure 5D). This interpretation was supported by analysis of individual gene loci, e.g. *CDKN1A* and *BBC3* (Figure 5E, F). As noted above, the vast majority of the synergy up p53 target genes also bind p53 in both nutlin-3 and combination-treated cells (Figure 4M, N and Figure 5D). In other words, synergy up p53 target genes, tend to bind p53 and BRD4 in nutlin-3-treated cells but p53 only in combination-treated cells (Figure 5D-F). These results suggest that in AML cells, BRD4 may act as a repressor of p53 target genes. In line with this idea, treatment of OCI-AML3 cells with CPI203 modestly activated a number of p53 target genes, even in the absence of nutlin-3, e.g. *PMAIP1, RAP2B, FUCA1* and others (Figure 3F). To directly test the ability of BRD4 to repress p53 target genes, we ectopically expressed the short form of BRD4 (BRD4S) in AML3 cells treated with nutlin-3 to stabilize p53. Cells ectopically expressing BRD4S activated p53 target genes *CDKN1A* and *PUMA* to a lesser extent than empty vector cells upon treatment with nutlin-3 (Figure 5G-I). Measured against a number of other p53 target genes, there was an invariable trend towards repression by BRD4S (Supplementary Figure 7A-H). The repressive effect of BRD4S was confirmed using a stably integrated p53 fluorescent/luminescent reporter gene (Supplementary Figure 6I, J). BRD4S also repressed p53 target gene expression in wild-type *TP53* MOLM13 cells (Supplementary Figure 7K-N). These results suggest that in AML, BRD4 is able to bind p53 target genes and can repress their activation, even when p53 is bound. Displacement of BRD4 by BET inhibition relieves this repression, leaving p53 free to activate its pro-apoptotic targets, thereby accounting for the synergistic killing of AML by combined MDM2 and BET inhibition.

## Discussion

In this study, we show that BRD4 inhibition together with p53 activation induces synergistic anti-leukemia activity both *in vitro* and *in vivo*, in AML retaining wild-type *TP53*. Synergy depends on wild-type *TP53*, is linked to enhanced activation of p53 by BETi and depends on expression of at least some p53 target genes, namely *BBC3* and *NOXA*. Regarding the mechanism underlying this synergy, we eliminated some obvious candidates, such as binding of p53 to its target genes and stabilization of p53 target mRNAs. Instead, we present evidence that the basis of this synergy is via relief of an unexpected repressive effect of BRD4 on expression of p53 target genes, thereby unleashing the full pro-apoptotic activity of p53 stabilized by MDM2i.

Through its role as a “reader” of acetylated lysine residues, BRD4 is widely regarded as a driver of transcriptional activation. Mechanistically, BRD4 promotes gene expression via the recruitment and activation of P-TEFb, which drives RNA polymerase II-dependent transcription (Jang et al., 2005). However, some reports have implicated BRD4 in transcriptional repression. Although some reports have implicated BRD4S preferentially as the repressive isoform, for example of transcriptionally silent latent HIV virus by recruitment of repressive SWI/SNF chromatin remodeling complexes (Conrad et al., 2017), other studies have shown that BRD4L also acts as a repressor, for example of the HPV-encoded E6 gene and autophagy genes by recruitment of histone methyltransferase G9A (Sakamaki et al., 2017; Wu et al., 2006). Conceivably, a BRD4/G9a interaction is also involved in repression of p53 target genes. Previous studies have demonstrated a physical interaction between p53 and BRD4 (Wu et al. 2013; Stewart et al. 2013). Wu et al showed that BRD4 promotes expression of p53 target gene, *CDKN1A*, at least in 293 cells (Wu et al., 2013). These studies raise the possibility that in some contexts a p53/BRD4 physical interaction promotes expression of p53 target genes, but in AML cells BRD4’s interaction with p53 is co-opted into a repressive interaction that silences p53 target genes and facilitates leukemogenesis. Our demonstration that BRD4 acts as a repressor and silences p53 activity extends the repressive activity of BRD4 into an important new context and has important implications for AML pathogenesis and candidate therapeutic approaches.

Regarding pathogenesis, across all human cancers, the *TP53* tumor suppressor gene is the most frequently mutated gene (Kastenhuber and Lowe, 2017). In mice, inactivation of p53 cooperates with activated RAS in leukemogenesis (Chan and Gilliland, 2004; Zhang et al., 2017; Zhao et al., 2010). However, although AML is, overall, a heterogeneous disease, surprisingly more than 90% of *de novo* AML retain wild-type *TP53* (Klco et al., 2013; Papaemmanuil et al., 2016), suggesting that human AML subtypes employ alternative mechanisms to inactivate the p53 pathway (Prokocimer et al., 2017). At least some AML dysregulate known p53 regulators, MDM2, MDM4 and ARF (Prokocimer et al., 2017). Our data suggest that in some AML dysregulation of BRD4 might also antagonize the p53 pathway to facilitate leukemogenesis. Consistent with this idea, *BRD4* exhibits elevated expression in ∼7% of AML (Bansal et al., 2017), and, in the majority of cases, this is accompanied by wild-type *TP53* (cBioportal).

With respect to novel candidate therapies for AML, our studies suggest promising efficacy of the MDM2i and BETi combination in AML. Previous studies have suggested that the efficacy of an MDM2i and BETi combination in chronic myeloid leukemia (CML) comes from dual targeting of the p53 and c-MYC pathways, by MDM2i and BETi respectively (Abraham et al., 2016). As a secondary consequence of BETi-mediated downregulation of c-MYC, BETi can activate at least one p53 target gene, *CDKN1A*, through loss of c-MYC-mediated gene repression (Mertz et al., 2011). Thus, although our studies indicate that *TP53* wild type AML are typically sensitive to the MDM2i/BETi combination, the dominant mechanisms underlying synergistic toxicity are likely to vary, especially given the genetic heterogeneity of AML (Klco et al., 2013; Papaemmanuil et al., 2016). At least in OCI-AML3 cells (representative of a recurrent AML genotype found in 10-15% of AML, *NPM1* and *DNMT3A* mutant and *TP53* wild type), we have shown an unexpected ability of BETi to directly potentiate activation of p53 by MDM2i, via relief of BRD4-mediated gene repression. Also in two mouse models, we observed markedly enhanced anti-AML activity of the drug combination, compared to either single drug alone. Of note here however, we used a clinical grade MDM2i, RG7112, that has been optimized against human MDM2i. Therefore, it is possible that our mouse studies diminish the on-target toxicity of MDM2i toward normal mouse tissues, thereby increasing the tolerability of the drug. Further validation of this novel drug combination, in terms of efficacy and tolerability, in both animal models and human cells is warranted.

The molecular heterogeneity of AML has been a hurdle to development of novel therapies of benefit to a substantial proportion of patients. For example, *NPM1*, the most commonly mutated gene in AML, is mutated in only 28% of AML and many genes are recurrently mutated, but in less than 10% of patients, for example, *ASXL1, IDH2, RUNX1* and *SRSF2* (Klco et al., 2013; Papaemmanuil et al., 2016). In contrast, in pre-clinical studies, a substantial proportion of AMLs are relatively sensitive to BETi and ∼90% of AML express wild-type *TP53* (Dawson et al., 2011; Filippakopoulos et al., 2010; Zuber et al., 2011)(Klco et al., 2013; Papaemmanuil et al., 2016)(Mertz et al., 2011), suggesting that the MDM2i and BETi drug combination can potentially target a majority of AML.

In summary, we demonstrate that MDM2 and BET inhibition are synthetically lethal in AML with wild-type p53. We propose that BRD4 represses transcription of p53 target genes, such that when BETi block repression in the context of p53 stabilization by MDM2i the p53 pathway is potently activated leading to synergistic killing in AML. As single agents these compounds have been shown to be effective and quite well tolerated in clinical trials (Alqahtani et al., 2019). Given the superiority of the combination over single drugs in our *in vitro* and *in vivo* studies, a clinical trial employing these two agents as a combination in AML retaining wild-type TP53 is justified.

## Acknowledgements

We thank Robert J. Sims, III and Jennifer A Mertz of Constellation Pharmaceuticals Inc. for providing CPI0610 and advice on *in vivo* studies and the staff of BICR Biological Services Unit for assistance with mouse experiments. Work in the lab of PDA was funded by CRUK program grant C10652/A16566. We thank support from Children with Cancer UK and the Howat Foundation (to KK^5^ and JC). KK^1^ was funded by Wellcome Trust (grant number 105641/Z/14/Z). Additional funding from CRUK Glasgow Centre (A25142) and Core Services at the Cancer Research UK Beatson Institute (A17196). This study was supported by the Glasgow Experimental Cancer Medicine Centre, which is funded by Cancer Research UK and the Chief Scientist’s Office, Scotland. Experiments in the lab of TC were supported by Sussex Cancer Fund. XH is a John Goldman Fellow [Leuka 2016/JGF/0005]

## Author contributions

Contributions to the manuscript: A-LL, PDA and MC conceived the idea for the project; A-LL and AN conducted most of the experiments; SL performed additional *in vitro* experiments; MT, LRD, JC, and CR assisted with *in vivo* experiments; HS and TC performed and supervised, respectively, experiments with primary human AML blasts; LM performed cell viability assays to calculate combination indices; KK^1^, JL, ST and MC provided technical assistance or advice; J-IS and KMR gifted reagents and shared critical unpublished data; WC performed RNA- and ChIP-sequencing; CM performed sequencing of TP53 in human AML blasts; KG, NR, XL and JC conducted bioinformatic analyses; XH assisted and advised on the MLL-AF9 mouse model; SA and TH assisted in procuring drugs for *in vivo* experiments; BXH gifted RG7112; KK^5^ and KB advised and supervised *in vivo* work; A-LL, AN, SL and PDA wrote the MS. PDA supervised the project. All authors have reviewed and approved the manuscript.

## Data Availability Statement

Chip Seq data has been deposited in the Gene Expression Omnibus (GEO) under accession codes GSE132246 and GSE132247. All other data supporting supporting the findings of this study are available from the corresponding author upon reasonable request.

## Methods

### Primary AML cells

A total of 15 AML patient samples were studied. All primary bone marrow aspirates were taken from routine diagnostic specimens after informed consent of the patients. The project received approval from the local ethics committee (The Brighton Blood Disorder Study, references 09/025/CHE and 09/H1107/1) and was conducted in accordance with the Declaration of Helsinki. Mononuclear cells from patients diagnosed with AML were isolated by Histopaque 1077 density gradient purification. They were stored in the Brighton and Sussex Medical School (BSMS) tissue bank and plated at a density of 40,000 cells/per well in the black 96 wells plates. Cells were plated in 80μl of RPMI containing 10% FCS, 100mM glutamine, 10,000 I.U/mL Penicillin and Streptomycin.

### AML Cell lines

THP1 was obtained from ATCC, KASUMI-1, MOLM-13 and OCI-AML3 from DSMZ, KG1a and MV411 were gifted from Professor Mhairi Copland and Dr. Xu Huang, respectively (both Paul O’Gorman Leukemia Research Centre, Glasgow). The authenticity of all cell lines was confirmed by genotyping. Cell lines were grown according to the vendors’ instructions, were incubated at 37°C in a humidified incubator with 5% CO_2_ and passaged every 3-4 days. For p53 activity assays with luciferase and GFP reporters, AML3 cells were stably infected with pGF-p53-mCMV-EF1α-Puro lentivirus (https://systembio.com/shop/pgf-p53-mcmv-ef1α-puro-ht1080-stable-cell-line/).

### Mice

Trib2-expressing AML blasts (Keeshan et al., 2006) from serial transplants were thawed and maintained in medium (DMEM with 15% FBS + 100 units/ml penicillin streptomycin and 2mM L-glutamine) and supplemented with 10ng/ml IL-3 (Peprotech, 213-13), 10ng/ml IL-6 (Peprotech 216-1) and 100ng/ml SCF (Peprotech 250-03). MLL-AF9 cells (Somervaille and Cleary, 2006) were maintained in medium (RPMI with 20% FBS + 100 units/ml penicillin streptomycin and 2mM L-glutamine) and supplemented with 10ng/ml IL-3 (Peprotech, 213-13). For *in vivo* experiments, mice were sub-lethally irradiated at 5.5Gy, and 4 hours later injected with 850,000 cells for the Trib2 murine AML model and 200,000 leukemia cells for the MLL-AF9 murine experiment. For both experiments a final volume of 200μl of leukemia cells was injected through the tail vein. Mice were maintained on Baytril antibiotic in their drinking water pre- and post-transplantation, for two weeks. Mice were bled via tail vein weekly to test for disease engraftment; when this was confirmed drug treatment was initiated. For *in vivo* studies, CPI0610 (Constellation Pharmaceuticals) was dissolved in heated 0.5% methylcellulose then sonicated (using a Diagneode Bioruptor). RG7112 was mixed with its vehicle supplied by Roche (2% hydroxypropyl cellulose, 0.1% polysorbate 80, 0.09% methyl paraben, 0.01% propyl paraben). For the Trib2 fixed end point experiment, Trib2 mice were treated for 21 days. RG7112 was initially given once daily 100mg/kg and CPI0610 twice daily 30mg/kg, both by oral gavage. However, due to excessive weight loss of one of the combination-treated mice, the mice received a 2 day “drug holiday” after 10 days and resumed dosing at once daily RG7112 70mg/kg and twice daily CPI0610 30mg/kg. For the Trib2 and MLL-AF9 survival experiments, mice initiated 21 days of drug dosing (not including 2 drug-free days every 5 days) (RG7112 was given once daily 70mg/kg and CPI0610 was given twice daily 30mg/kg, both by oral gavage). After reaching the end of treatment, mice were culled immediately or maintained to an ethical survival end point.

### Drug treatments of cells *in vitro*

For drug treatment of primary AML blasts, 2.5μM nutlin-3 alone, 200nM CPI203 alone, and the two in combination were added to triplicate wells to a volume of 100ul. Control experiments were performed without the addition of drugs using vehicle (DMSO) only. After 48hrs cell viability was measured using the CellTiter-Glo reagent (Promega, G7572), and the luminescence was detected with Biotek synergy HT plate reader and analyzed using Gen 5 version 1.08 software. Cell viability was also determined by trypan blue dye exclusion in some experiments (Sigma, T8154). Apoptosis was assessed used Annexin V and propidium iodide (PI) staining (BD Biosciences; 556547) as per manufacturer’s instructions and analyzed using the BD Fortessa flow cytometer. For p53-promoter based dual GFP/Luminescence assay after 24h 2.5 M nultin-3 treatment, cells were lysed using 1xpassive buffer (Promega, E1941) and lysate was either measured directly for GFP signal or incubated with luciferase assay reagent LAR (Promega, E1500) for luminescence signal. The GFP and luminescence were detected with CLARIOstar plate reader and analyzed using Microsoft Excel.

### Drug combination and synergy analyses

Chou-Talalay Combination Index method: AML cell lines were treated with combinations of CPI203 800nM and nutlin-3 ranging from 1.25μM to 10μM) to determine synergistic doses. Cell viability was determined after 72 hours by either trypan blue dye (Sigma, T8154) exclusion or resazurin (Alamar blue dye, Sigma) with the Envision Fluorescent Reader (Perkin Elmer). Mean fluorescent values from multiple replicates were calculated for each condition. Assessment of synergy was made by calculating combination indices (CI) using Calcusyn software (version 2.0), CI < 1 considered synergistic, CI=1 considered additive and CI>1 considered antagonistic (Chou and Talalay, 1984). Bliss Independence method: AML cell lines were seeded on 384-well plates at 3,000 cells per well and treated for 72 hours (or 24 hours for BET PROTAC experiments) with the indicated doses (2x serial dilution) of drugs, alone or in combination. Luminescence from quadruplicate was measured using the CellTiter-Glo reagent (Promega, G7572). Bliss synergy was calculated using SynergyFinder v2.0 web-based application with default parameters for calculating bliss independence. The difference between the observed combined effect and the expected combined effect of the two drugs is called the Excess over Bliss (*eob*). Positive *eob* values are indicative of synergistic interaction, negative *eob* values are indicative of antagonistic behavior and null *eob* values indicate no drug interaction.

### RNA extraction, cDNA synthesis and qPCR

RNA was extracted using the RNeasy Mini kit (Qiagen, cat no. 7410) according to the manufacturer’s protocol. On-column DNA digestion was carried out using DNase1 (Qiagen, cat no. 79254). Eluted RNA was quantified using the Nanodrop 2000 (ThermoFisher Scientific) and 1μg RNA was used in cDNA synthesis (Invitrogen, 18080-093), according to the manufacturer’s instructions. For cDNA synthesis: 1μg RNA, 1μl of 50 μM oligo DT primer (Invitrogen 18418020), 1μl dNTP mix (10mM each of dATP, dGTP, dCTP, and dTTP) and water to a final volume of 14μl were mixed and heated at 65°C for 5 minutes. After a 1-minute incubation on ice, the following were added: 4μl 5X first strand buffer, 1μl 0.1M DTT, 1μl Superscript™ Reverse Transcriptase (Invitrogen, 18080-093), and heated at 25°C for 5 minutes. Tubes were then heated to 55°C for 1 hour and the reaction was inactivated by a 15-minute incubation at 70°C. qPCR was done using 10μl of SYBR green master mix (2X DyNAmo HS SYBR green qPCR master mix, Thermo-Scientific, F-410), 200nM of each primer of interest and water was added to a final volume of 20μl as a master mix to 1μl of DNA in Hard Shell PCR Plates, 96-well white, Bio-Rad, (HSP9601). The primer sequences used are listed in Supplementary Table 4. The PCR reactions were performed as follows; 95°C for 3 minutes, 95°C for 10 seconds, 60°C for 20seconds, 72°C for 30 seconds, steps 2 to 4 repeated 39 times, 72°C for 5 minutes, and 65°C for 5 seconds then a gradient up to 95°C for melt curve analysis.

### mRNA stability assays

OCI-AML3 cells were treated with Vehicle (DMSO), 200nM CPI203, 2.5μM nutlin-3 or the drug combination. After a two-hour incubation, cells were treated with 5μM Actinomycin D and harvested after 0, 2, 4 and 6 hours of Actinomycin D treatment. Cell pellets were stored overnight at −80°C before RNA was extracted and cDNA synthesized for qPCR analysis as described above. Abundance of *CDKN1A* and *BBC3* mRNA at each time point was calculated relative to time 0 using the delta CT method (Ct at time 0 minus Ct of each time point).

### RNA sequencing

Sequencing libraries were made from poly-A RNA, as recommended by Illumina, and sequenced using either an Illumina GAIIX or a NextSeq 500 sequencer. RNA-seq paired-end reads were assessed for quality using the ‘FastQC’ algorithm, trimmed as appropriate using the algorithm ‘trim-galore’ (version 3.0), then aligned to the human genome using the splice-aware aligner TopHat2. Reference splice junctions were provided by a reference transcriptome from the Ensembl GRCh37 (hg19) build, release version 73. The Cuffdiff tool from the Cufflinks suite was used to process aligned reads and perform maximum likelihood estimation to assess transcript abundances, before calculating the differential expression of transcripts across samples. In parallel, aligned reads for genic isoforms were collated and total read counts per gene were calculated using htseq-count version 0.5.4p3, before differential expression analysis using the linear modelling tool DESeq2 was performed. Using both differential expression methods, significantly changing expression was defined as an FDR-corrected p-value ≤0.005. FPKM (Fragments Per Kilobase of transcript per Million mapped reads) values were then generated. Gene ontology analysis was performed using Gene Set enrichment Analysis (GSEA), DAVID (version 6.7) and Ingenuity^®^ Pathway Analysis (IPA) software.

### ChIP-sequencing

ChIP-seq protocol was adapted from previously (Kirschner et al., 2015; Schmidt et al., 2009). Antibody-bound magnetic beads (anti-rabbit or anti-mouse, as appropriate) (Dynabeads beads, M280) were incubated with 10μg of BRD4 antibody (Abcam, 128874) or 10μg of p53 antibody (Santa Cruz, DO1) (rabbit or mouse IgG was used as a negative control (Sigma, M7023)) for 4 hours at room temperature, then added to the lysed samples. During the 4-hour antibody-bead incubation, 20×10^6^ OCI-AML3 cells were treated with vehicle, 200nM CPI203, 2.5μM nutlin-3 or the drug combination for 6 hours. Following this, cells were pelleted at 200g for 5 minutes and re-suspended in serum free RPMI media and cross-linked for 10 minutes on a rocker by adding 16% methanol-free paraformaldehyde (Alfa Aesar, 43368) (final concentration 1%). Cross-linking was quenched with 2.5M glycine (final concentration 0.125M) for 5 minutes. Cells were washed twice in cold PBS. Three lysis buffers (LB1-3) were used to produce chromatin lysates and the following protease inhibitors were added to each: 1x protease inhibitor cocktail (Sigma, P8340) and 50μg/ml PMSF (Sigma, P7626). 10ml of lysis buffer 1 (LB1) (50mM Hepes-KOH, pH 7.5; 140mM NaCl; 1mM EDTA; 10% glycerol; 0.5% NP40 and 0.25% Triton X-100) was added to the pellet of cross-linked cells and this was rocked at 4°C for 10 minutes, then centrifuged at 2,000 x g for 4 minutes at 4°C. The supernatant was discarded, and the pellet then re-suspended in 10ml of LB2 (10mM Tris-HCl, pH 8.0; 200nM NaCl; 1mM EDTA; 0.5mM EGTA), and rocked at 4°C for 5 minutes. The suspension was again pelleted, the supernatant discarded and the pellet then re-suspended in 2ml of LB3 (10mM Tris-HCl, pH 8; 100mM NaCl; 1mM EDTA; 0.5mM EGTA; 0.1% Na-deoxycholate; 0.5% N-lauroylsarcosine). This 2ml lysate was transferred to a FACS tube suitable for sonication. A Soniprep 150 set at 14 microns amplitude was used for 12 cycles of sonication (each cycle was 30 seconds on, 60 seconds off). Sonication was performed one sample at a time on ice. Following sonication, Triton-X-100 was added to the lysates (final concentration 1%), and lysates were centrifuged at 16,000 x g for 12 minutes at 4°C to pellet the debris. 50μl of the sonicated lysate from each treatment condition was stored at −20°C as input control. Cell lysates were pre-cleared with 100μl of magnetic beads. Following this, the 100μl of antibody-bead mix was then added to the cell lysates for overnight immunoprecipitation on a rotating platform at 4°C. Following completion of this incubation, the beads were washed 5 times in 1ml of RIPA buffer (50mM Hepes-KOH, pH 7.5; 500mM LiCl; 1mM EDTA; 1% NP-40; 0.7% Na deoxycholate plus protease inhibitors as described above). The pellets were then washed in 1ml of TBS (20mM Tris-HCl, pH 7.6; 150mM NaCl), centrifuged at 960 x g for 3 minutes at 4°C and then, following removal of the supernatant, the ChIP samples were eluted off the magnetic beads with 200μl of elution buffer (50mM Tris-HCl, pH 8; 10mM EDTA; 1% SDS) and reverse cross-linked by heating at 65°C for 6 – 18 hours. The beads were vortexed every 5 minutes for the first 15 minutes. 200μl of TE was added to each IP and Input samples. 8μl of 1mg/ml RNaseA (Ambion, 2271) was added and the samples incubated at 37°C for 30 minutes. 4μl of proteinase K (20mg/ml) (Invitrogen, 25530-049) was then added and the samples incubated at 55°C for 1-2 hours. The samples were then purified using the Qiagen MinElute PCR purification kit (28006), eluted in 30μl elution buffer and stored at −20°C until required for qPCR or sequencing.

ChIP-seq libraries were made as recommended by Illumina, using the NEBNext Ultra DNA Lib Prep kit (E7370S) and NEBNext Multiplex Oligos, (Index Primers Set 2; E7500S). The quality of the libraries was assessed using the Agilent 2200 tape station (Agilent Technologies), with the Agilent D1000 ScreenTape system (Agilent, 5067-5582) and D1000 reagents (Agilent, 5067-5583). Sequencing was performed on an Illumina NextSeq 500 sequencer using the High Output kit v2 to perform 75 cycles of single-read sequencing.

Sequenced single-end ChIP-seq reads were trimmed using trim-galore (version 3.0) and aligned to the hg19 build of the human genome using Bowtie2 alignment software. Reads with a Phred quality score of <15 were removed. Non-unique or duplicate reads were removed using samtools and Picard tools (v1.98) respectively. Regions of BRD4 occupancy were determined using the SICER algorithm with a redundancy threshold of 1, a window size of 200, a fragment size of 150, an effective genome fraction of 0.75, a gap size of 200 and an FDR corrected p-value of 0.01 (suited to broader BRD4 peaks). To reduce experimental bias, a robust peak set, defined as being present in both replicates using the bedtools intersect function, was used. Regions of p53 occupancy were determined using the Macs tool (version 1.4) with a p-value cutoff of 1e-5 (suited to narrower peaks), then robust peaks that were present in at least 2 replicates were identified using the bedtools intersect function. BRD4 and p53 peaks were visualized using either the UCSC web interface or the WashU browser.

### Ectopic expression and gene knockdown by shRNA or CRISPR

Gene knockdown or overexpression in OCI-AML3 cells was achieved by viral transfection. DNA constructs were used to generate virus for infection of OCI-AML3 cells to generate knock-down or over expression (pTRIPZ and pLKO.1 for knock-down, pLentiCRISPR V2 for CRISPR knock-out, and pLenti4 and pCDH for over expression (Supplementary Table 5)). To generate virus, 2×10_6_ HEK-293T (ATCC, CRL-11268) or HEK 293FT (gifted from Dr David Bryant, Beatson Institute for CRUK) were plated one day prior to transfection at 2 million cells per plasmid in a 10 cm plate in culturing media (DMEM with 10% v/v FBS and L-glutamine 2mM and penicillin streptomycin 100 units per ml). Transfection was carried out in antibiotic-free culturing media (6ml) using 10μl lipofectamine (Lipofectamine ® 2000Reagent, 1mg/ml, Invitrogen P/N 2887) in 400μl Opti-mem media (Gibco Opti-mem reduced serum media, Life Technologies 31985062). This was combined with a further 400μl Opti-mem containing 2.5 μg of plasmid DNA, 1.86μg of ps PAX2 and 1μg of VSVG, incubated for 20-25 minutes at room temperature and then added dropwise onto the target plate of HEK-293T. After 6 hours the medium was aspirated and re-freshed with 6mls of normal culture medium containing antibiotic as above. Two days after transfection of the HEK-293T, infection of the target cells was initiated by spinning down 1.5×10^6^ target cells per sample and re-suspending in 400μl of culture media in a 12-well plate. 6ml of virus-containing media was removed from the plate of 293T cells (the 293T cells were refreshed with 6mls of antibiotic-containing media to allow a second infection the following day), filtered through a 0.45μM filter and mixed with polybrene (8μg/ml (Millipore TR-1003). To each well of target cells, 1ml of virus-containing media was added. A GFP-expressing virus and a ‘cells only’ well were included as controls for the transfection and infection respectively. The 12-well plate was then spun at 2500rpm at room temperature for 90mins and incubated at 37°C for 3-6hrs. Cells were then transferred to a 25cm flask with 9mls of media for an overnight incubation. The following day a second infection was carried out as described above, and 24 hours after this the cells were selected with 1μg/ml puromycin.

### Immunoblotting and antibodies

Cells were lysed directly in 1x Laemmli sample buffer (2% SDS, 10% glycerol, 0.01% bromophenol blue, 62.5mM Tris, pH 6.8) and boiled for 4 minutes. 30μg of protein was loaded into each lane of the gel. The polyacrylamide gels (BioLegend 456-1095 or NOVEX) were run at 150V in 1x running buffer (25mM Tris, 192mM glycine, 0.1% SDS, pH8.3). Proteins were transferred onto PVDF membranes in transfer buffer (20% methanol, 25mM Tris, 192mM glycine, 0.01% SDS, 20%) for 1 hour 25 minutes at 60V at room temperature. Membranes were dried on top of filter paper and reactivated with methanol, blocked in TBS with 5% milk or 4% BSA and probed with antibodies overnight at 4°C. β-Actin was used as a loading control for all western blots (1 in 200,000 dilution, Sigma A1978). All antibodies were diluted in 5% BSA TBS pH 7.5 with 0.05% sodium azide. Primary antibodies to the following targets were used at 0.1-1μg/ml: Actin (Sigma, A1978, RRID:AB_476692), CDKN1A (Santa Cruz, 397, RRID:AB_632126), c-MYC (Santa Cruz, 764, RRID:AB_631276), MDM2 (Santa Cruz, 965, RRID:AB_627920), PARP (Cell Signalling, 9542, RRID:AB_2160739), BBC3 (Imgenex, 458, RRID:AB_1151450), TP53 (DO1) (Santa Cruz, 126, RRID:AB_628082), NOXA (Abcam, 13654, RRID:AB_300536), BRD4 (Abcam, 128874, RRID:AB_11145462), BCL-2 (Cell signaling, 2870, RRID:AB_2290370) (Supplementary Table 6). The following morning membranes were washed in TBS pH 7.5 (3 x 5 minute washes) and then incubated with species-matched HRP-linked secondary antibody. The following secondary antibodies were added for a 1-hour incubation at room temperature: anti-mouse IgG HRP-linked (Dako, p0447) and anti-rabbit IgG HRP-linked (Cell signaling, 7074s), both at a dilution of 1:5,000. Following incubation with secondary antibodies, membranes were given 3 x 10-minute washes in TBS pH7.5 + 0.1%Tween 20 and then incubated in ECL Western blotting chemiluminescent substrate (Thermo Scientific, 32106) for 1 minute. Visualisation of protein bands was then carried out using either GE Healthcare Amersham hyperfilm on a Kodak X-Omat 480 RA X-ray processor, or BIORAD ChemiDoc Imaging system.

### FACS analysis

For flow cytometric evaluation of apoptosis in AML cell lines and primary murine AML cells treated with the above single agents or combination treatments, apoptosis was measured by combined Annexin-V/propidium iodide (PI) staining (Cambridge Bioscience, K101-400), myeloid differentiation was assessed using stains for CD11b (eBioscience, 12-0081) and GR1 (eBioscience, 25-5931), lymphoid differentiation was assessed using stains for CD4 (Biolegend, 100553), CD8a (eBioscience, 12-0081), CD19 (eBioscience, 47-0193080) and B220 (eBioScience, 45-0452). Drug or vehicle-treated cells were collected and first washed with 1ml of PBS. Cells were then re-suspended in 1ml of 1X Binding Buffer. Following this, the relevant stain of interest was added and samples incubated for 10 minutes at room temperature in the dark. Samples of human cell lines were then analyzed on a FACSCalibur (BD Biosciences) and primary murine samples were analyzed on a FACSCanto (BD Bioscience).

Cells were centrifuged at 1000rpm for 5 minutes and resuspended in 1ml sorting buffer (PBS 2 % FBS) and DAPI was added immediately prior to sort (final concentration 1μg/ml; 1 in 1000 dilution). Cells were sorted by Jennifer Cassells at the Paul O’ Gorman Leukaemia Research Centre using a BD FACSARIA III sorter and were collected in 1ml sorting buffer, pooled, centrifuged as above and left to recover in culturing medium at 37°C. Gating strategy for sorting GFP Positive cells was live cells>cell profile>single cells>GFP positive cells.

For assessment of disease engraftment and disease burden in mice, blood was collected via tail vein. Red blood cell lysis was carried out using BioLegend 10x red blood cell lysis buffer as per manufacturers’ instructions (incubation time was increased to 8 minutes after optimization). After centrifugation, cell pellets were resuspended in 400μl PBS and % GFP+ was measured using the BD Fortessa flow cytometer. At mouse cull, leg bones were cleaned upon removal then crushed using a pestle and mortar. The cell suspension was filtered and washed with ice-cold PBS+2% FBS. For spleen and thymus, the organ was placed in a 70μM cell strainer and pushed through using the plunger of a 5ml syringe into a 50ml falcon containing PBS+2% FBS. Suspensions were pelleted at 200g for 5 minutes, and the pellets resuspended in red cell lysis buffer before being pelleted as above. Finally, cells were washed in 1ml PBS+2% FBS. For FACS, compensation was calculated using ULTRACOMP beads (E biosciences 01-2222-41) incubated for 20 minutes with each antibody used in the experiment. Fluorescence minus one (FMO) controls were also included using cells incubated with a mix of all antibodies except one in each FMO control. A master mix was prepared containing all antibodies at a 1 in 250 dilution (myeloid cells [CD11b, E-bioscience, 17-0112-83; Gr1, E-bioscience, 25-5931-81], B cells [B220, E-bioscience, 45-0452-80; CD19, E-bioscience, 47-0193080], T cells [CD4, Biolegend, 100547; CD8, E-bioscience, 12-0081-82] (Supplementary Table 6)) in PBS+2% FBS and cell pellets were re-suspended in 50μl of master mix for a 30 minute incubation on ice and away from light. Immediately prior to FACS analysis DAPI was added at a 1 in 1000 dilution (1μg/ml) (Sigma, D9542-1MG).

### Statistical analysis

Results were statistically analyzed in GraphPad Prism (version 7.0, GraphPad Software Inc., San Diego, CA) and presented as mean±SEM. Statistically significant differences between two groups were assessed by two-tailed unpaired t-test. p≤0.05 was considered significant. ns, not significant; *p≤0.05; **p≤0.01; ***p≤0.001; ****p≤0.0001 for indicated comparisons. Statistical details of each experiment can be found in the Results and Figure Legend sections.

## Supplemental Information, Figure Titles and Legends

**Supp Figure 1.**
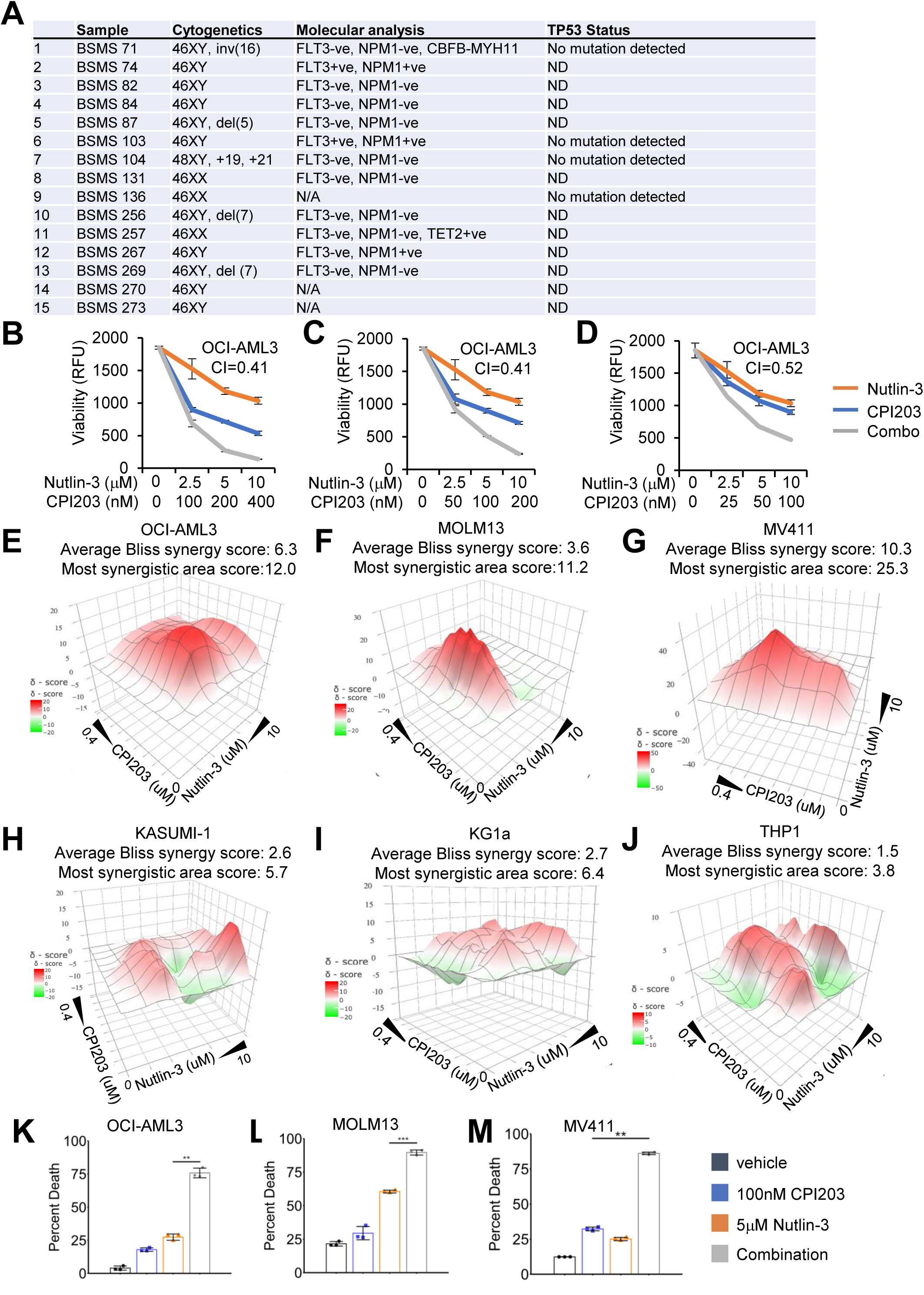
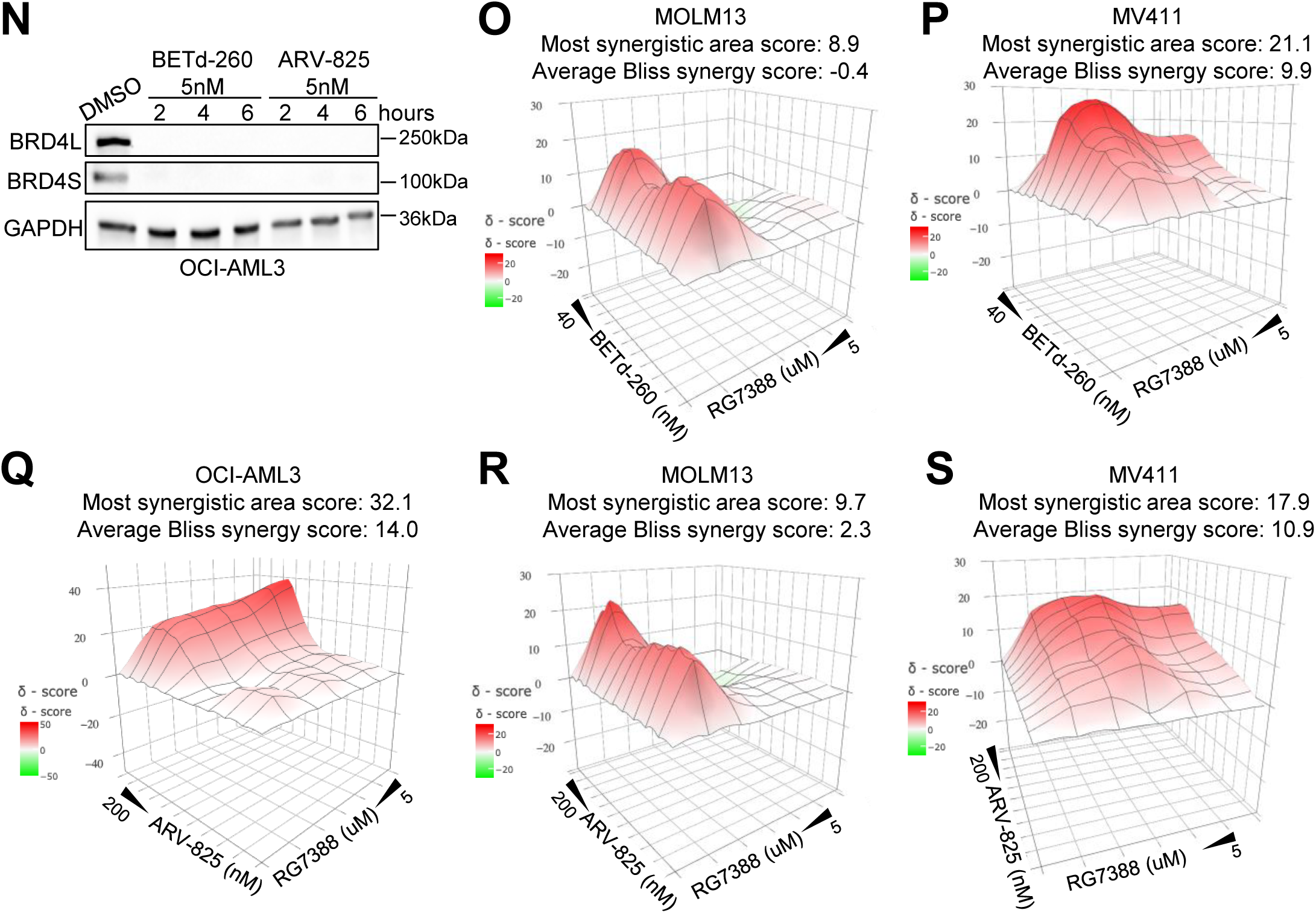
MDM2 and BET inhibitors are synergistically lethal to primary human AML blasts and AML cell lines with wild-type *TP53*. **A.** Summary of the cytogenetic, molecular features and *TP53* status of the 15 primary human AML samples treated with the drug combination *in vitro* in Figure 1A. FLT3+ve denotes mutation for FLT3 (FLT3-ITD is the recurrent mutation seen in AML) and FLT3-ve denotes WT. NPM1+ve denotes recurrent AML mutation. NMP1-ve denotes WT. For the 4 indicated samples, all exons of TP53 were sequenced, but no mutations detected. ND: not determined. N/A: not available. **B.** OCI-AML3 cell viability (each treatment in triplicate) was assessed by resazurin assay after 72 hours, using a treatment ratio of CPI203:nutlin-3 of 1:25 (Means +/− SD are shown, n=3). **C.** OCI-AML3 cell viability (each treatment in triplicate) was assessed by resazurin assay after 72 hours, using a treatment ratio of CPI203:nutlin-3 of 1:50 (Means +/− SD are shown, n=3). **D.** OCI-AML3 cell viability (each treatment in triplicate) was assessed by resazurin assay after 72 hours, using a treatment ratio of CPI203:nutlin-3 of 1:100 (Means +/− SD are shown, n=3). **E.** Excess Over Bliss plot showing synergistic effects between Nultin-3 and CPI203 in *TP53* wild type OCI-AML3 cells. Cell viability (each treatment in quadruplicate) was assessed by CellTiter-Glo after 72 hours. Bliss synergy score are indicated. **F.** Excess Over Bliss plot showing synergistic effects between Nultin-3 and CPI203 in *TP53* wild type MOLM13 cells. Cell viability (each treatment in quadruplicate) was assessed by CellTiter-Glo after 72 hours. Bliss synergy score are indicated. **G.** Excess Over Bliss plot showing synergistic effects between Nultin-3 and CPI203 in *TP53* wild type MV411 cells. Cell viability (each treatment in quadruplicate) was assessed by CellTiter-Glo after 72 hours. Bliss synergy score are indicated. **H.** Excess Over Bliss plot showing synergistic effects between Nultin-3 and CPI203 in *TP53* mutant KASUMI-1 cells. Cell viability (each treatment in quadruplicate) was assessed by CellTiter-Glo after 72 hours. Bliss synergy score are indicated. **I.** Excess Over Bliss plot showing synergistic effects between Nultin-3 and CPI203 in *TP53* mutant KG1a cells. Cell viability (each treatment in quadruplicate) was assessed by CellTiter-Glo after 72 hours. Bliss synergy score are indicated. **J.** Excess Over Bliss plot showing synergistic effects between Nultin-3 and CPI203 in *TP53* mutant THP1 cells. Cell viability (each treatment in quadruplicate) was assessed by CellTiter-Glo after 72 hours. Bliss synergy score are indicated. **K.** Cell kill in the OCI-AML3 cell line as assessed by flow cytometry using annexin-V and propidium iodide stains, according to treatment condition (72hrs) (***=p≤0.001). Percent death = sum of percent single (annexin-V+ or propidium iodide+) and double positive cells (two tailed unpaired t-test, Means +/− SD are shown, n=3). **L.** Cell kill in the MOLM13 cell line as assessed by flow cytometry using annexin-V and propidium iodide stains, according to treatment condition (72hrs) (***=p≤0.001). Percent death = sum of percent single (annexin-V+ or propidium iodide+) and double positive cells (two tailed unpaired t-test, Means +/− SD are shown, n=3). **M.** Cell kill in the MV411 cell line as assessed by flow cytometry using annexin-V and propidium iodide stains, according to treatment condition (72hrs) (**=p≤0.01). Percent death = sum of percent single (annexin-V+ or propidium iodide+) and double positive cells (two tailed unpaired t-test, Means +/− SD are shown, n=3). **N.** Western blot of BRD4 in OCI-AML3 cells treated with BETd-260 or ARV-825 at 5nM for different times. **O.** Excess Over Bliss plot showing synergistic effects between RG7388 and BETd-260 in MOLM13 cells. Cell viability (each treatment in quadruplicate) was assessed by CellTiter-Glo after 24 hours. Bliss synergy score are indicated. **P.** Excess Over Bliss plot showing synergistic effects between RG7388 and BETd-260 in MV411 cells. Cell viability (each treatment in quadruplicate) was assessed by CellTiter-Glo after 24 hours. Bliss synergy score are indicated. **Q.** Excess Over Bliss plot showing synergistic effects between RG7388 and ARV-825 in OCI-AML3 cells. Cell viability (each treatment in quadruplicate) was assessed by CellTiter-Glo after 24 hours. Bliss synergy score are indicated. **R.** Excess Over Bliss plot showing synergistic effects between RG7388 and ARV-825 in MOLM13 cells. Cell viability (each treatment in quadruplicate) was assessed by CellTiter-Glo after 24 hours. Bliss synergy score are indicated. **S.** Excess Over Bliss plot showing synergistic effects between RG7388 and ARV-825 in MV411 cells. Cell viability (each treatment in quadruplicate) was assessed by CellTiter-Glo after 24 hours. Bliss synergy score are indicated.

**Supp Figure 2.**
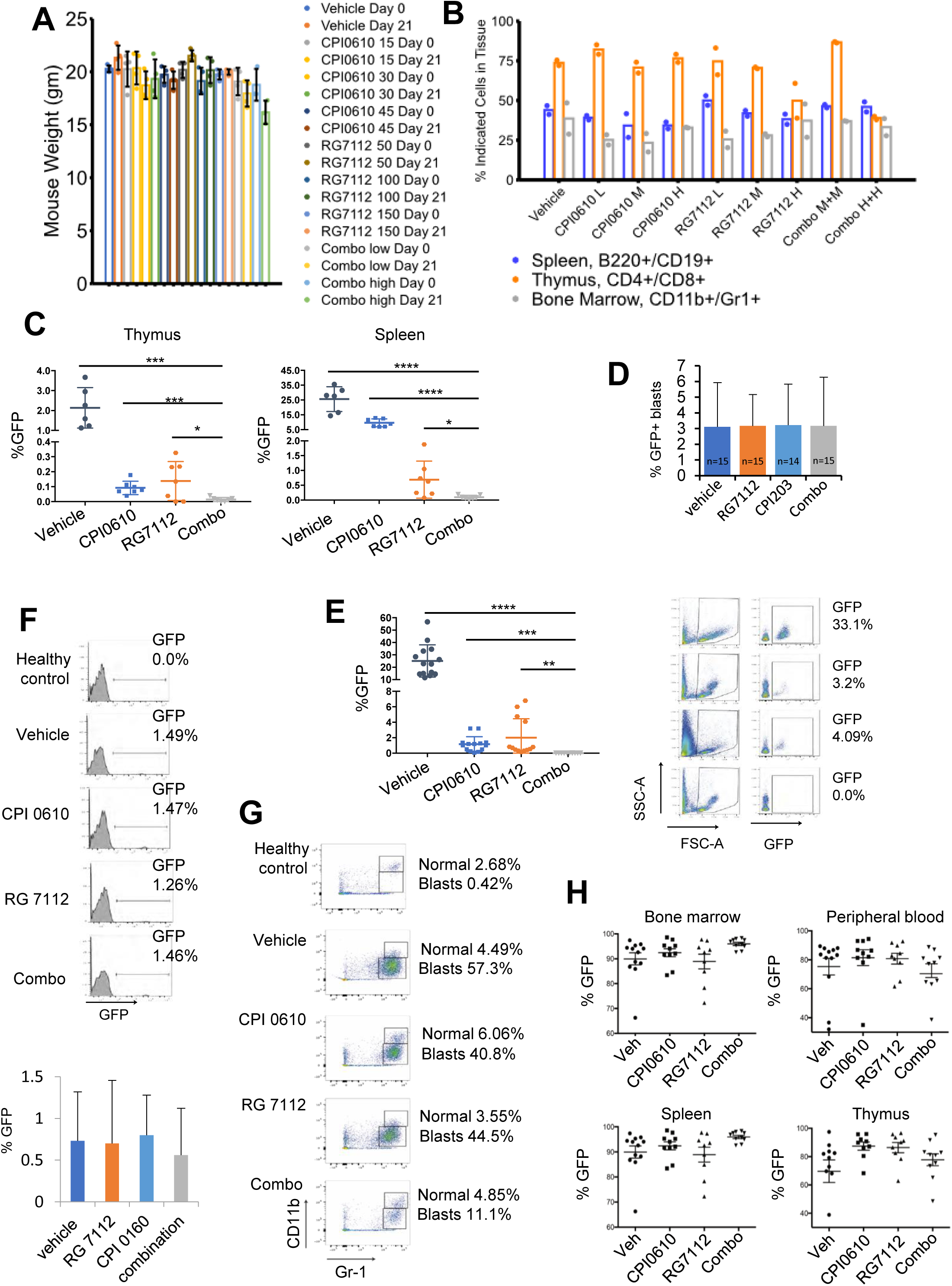
MDM2 and BET inhibitors cooperate to eradicate AML in *in vivo* mouse models. **A.** Normal non-leukemic mouse weights at indicated daily drug doses (mg/kg) at day 0 and on completion of drug treatment (day 21). For combo, low dose = 30mg/kg CPI0610 and 100mg/kg RG7112; high dose = 45mg/kg CPI0610 and 150mg/kg RG7112 (Means +/− SD are shown, n = 4 for all except for combo high day 0 and combo high day 21, where n = 3). **B.** Percent of indicated cell types in indicated tissue in normal non-leukemic mice treated with vehicle, low, medium or high dose of CPI0610 (15, 30 or 45mg/kg), RG7112 (50, 100, 150mg/kg) or combination (M+M = 30mg/kg CPI0610 and 100mg/kg RG7112; H+H = 45mg/kg CPI0610 and 150mg/kg RG7112). **C.** Disease burden (percent GFP+ blasts in thymus or spleen) in Trib2 mice at the end of 21 days of drug treatment (****=p≤0.0001, ***=p≤0.001, *=p≤0.05, by two tailed unpaired t-test, Means +/− SD are shown, n = 6 for vehicle, n=7 for single drug and combo conditions). **D.** Disease burden (percent GFP+ blasts in peripheral blood) in Trib2 mice pre-drug treatment (Error bars are SD, n for each group is labeled) **E.** Disease burden (percent GFP+ blasts in peripheral blood) in Trib2 treated mice at the end of 21 days treatment in peripheral blood according to treatment condition. Right panel shows representative FACS data (****=p≤0.0001, ***=p≤0.001, **=p≤0.01, two tailed unpaired t-test, Means +/− SD are shown, n=14, gating strategy for sorting GFP Positive cells was live cells>cell profile>single cells>GFP positive cells). **F.** MLL-AF9 disease burden (percent GFP+ blasts in peripheral blood) pre-drug treatment. Top shows representative FACS data (Gating strategy for sorting GFP Positive cells was live cells>cell profile>single cells>GFP positive cells.). **G.** Representative MLL-AF9 disease burden in peripheral blood, based on percent CD11b^low^ Gr1-expressing immature blasts known to expand in AML (Keeshan et al. 2006), after 7 days of drug treatment. Also shown are the more normal, mature myeloid cells (CD11b^high^ Gr1+) (Gating strategy for sorting GFP Positive cells was live cells>cell profile>single cells>GFP positive cells.). **H.** % MLL-AF9 GFP+ blasts at time of cull in indicated tissues after indicated drug treatments.

**Supp Figure 3.**
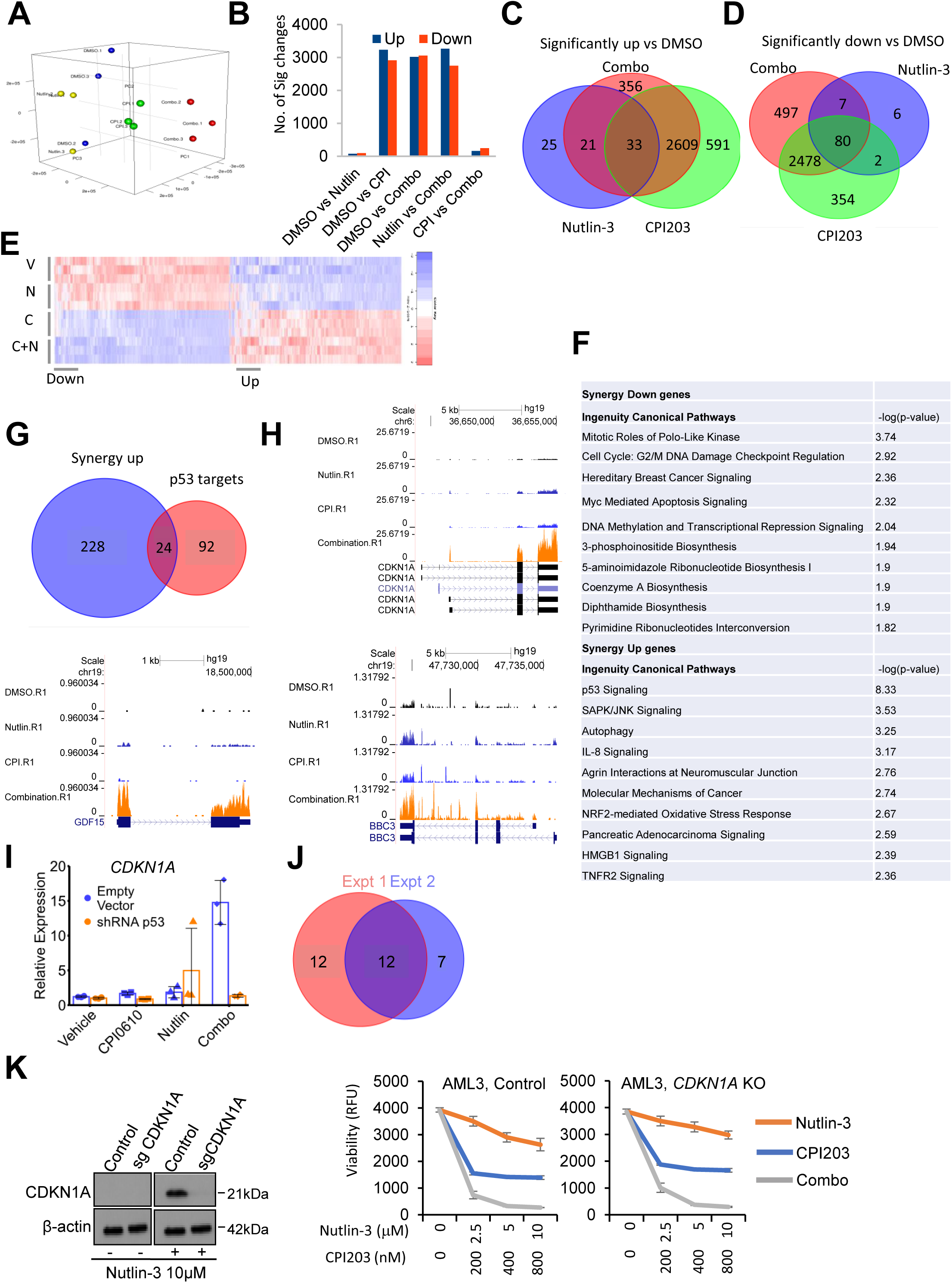
Figure 3. BET inhibitors potentiate activation of p53 target genes by p53. **A.** A principal component analysis of RNA-seq data of the OCI-AML3 cell line (from Figure 3E), according to treatment condition. **B.** Number of significant gene expression changes between treatment conditions in RNA-seq data of treated OCI-AML3 cells. **C.** Venn diagram of genes significantly up-regulated in RNA-seq data, according to treatment condition of OCI-AML3 cells. **D.** Venn diagram of genes significantly down-regulated in RNA-seq data, according to treatment condition of OCI-AML3 cells. **E.** Heat map of all significant changes in gene expression in RNA-seq data of OCI-AML3 cells according to treatment condition. **F.** An IPA analysis of synergistically up-regulated and down-regulated genes (synergy as defined in Figure 3E, p-values from fisher’s exact test). **G.** Venn diagram showing the number of high-confidence p53 target genes synergistically up-regulated by the drug combination. **H.** Representative examples of RNA-seq gene tracks for the p53 target genes *CDKN1A*, *GDF15* and *BBC3*, according to treatment condition. **I.** qPCR assessment of expression of CDKN1A in control (empty-vector) OCI-AML3 cells and shRNA p53 OCI-AML3 cells, according to treatment condition (Means +/− SD are shown, n=3). **J.** Venn diagram showing number of expression changes in p53 target genes in OCI-AML3 cells expressing wild-type p53 in the first and second RNA-seq experiments (i.e. Figures 3F and 3N, respectively). **K.** Western blot analysis of control (empty-vector) OCI-AML3 cells and CRISPR/CAS9 CDKN1A knock out OCI-AML3 cells, treated with vehicle or 10μM nutlin-3 (left); and a resazurin analysis of control (empty-vector) OCI-AML3 cells and CDKN1A knock out OCI-AML3 cells according to treatment condition, following 72 hours of drug treatment (right) (Means +/− SD are shown, n=3).

**Supp Figure 4.**
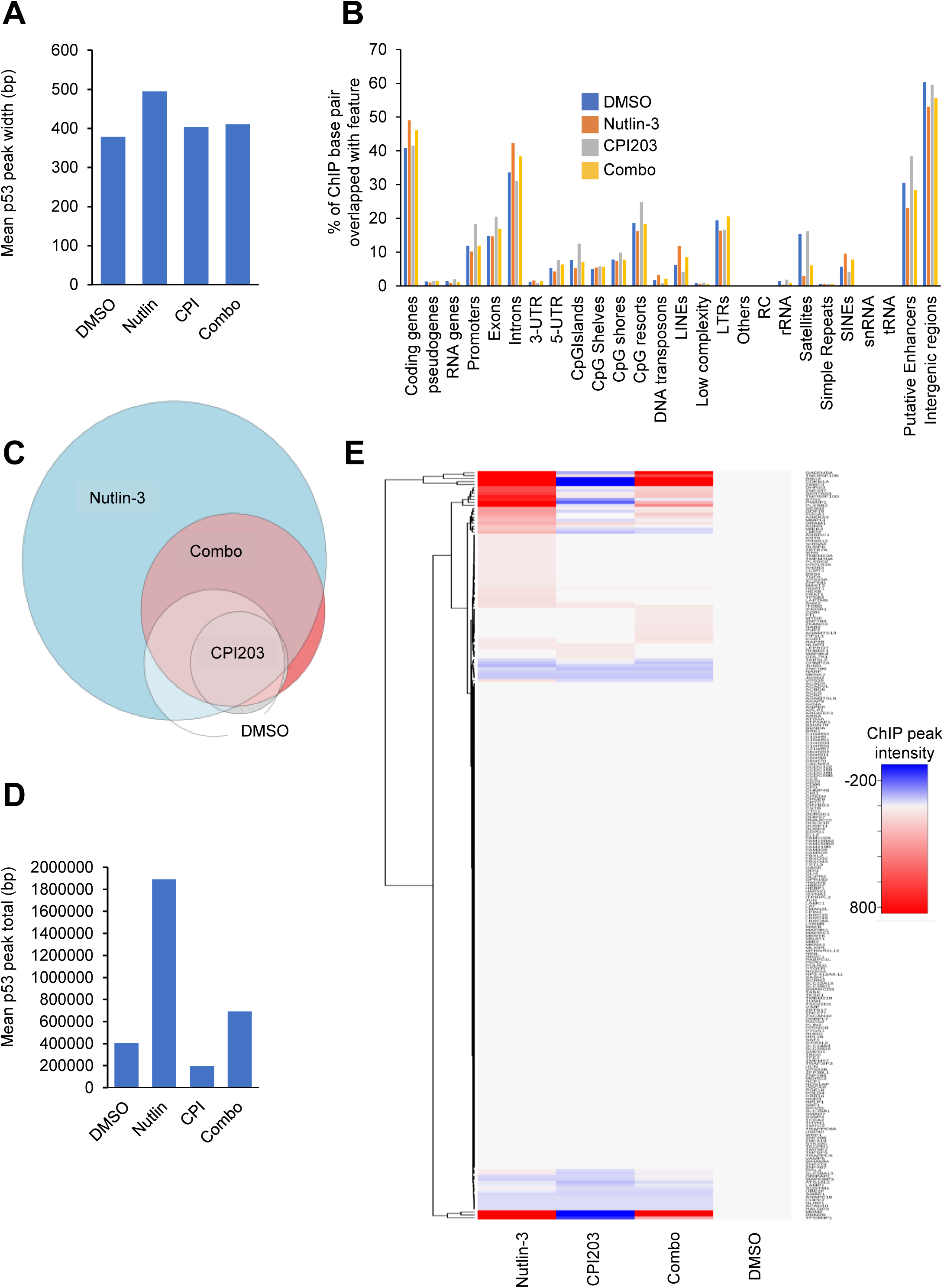
BET inhibitors do not stabilize p53 target mRNAs nor increase binding of p53 to target genes. **A.** Mean p53 peak width in base pairs (peaks present in at least 2 out of 3 ChIP-seq replicates; mean of 2- 3 replicates) after the indicated drug treatments of OCI-AML3 cells. **B.** Genomic distribution of p53 binding sites identified by ChIP-seq (peaks present in at least 2 out of 3 replicates) after the indicated drug treatments of OCI-AML3 cells. **C.** Venn diagram showing relative overlap of p53 ChIP-seq peaks (peaks present in at least 2 out of 3 replicates) after the indicated drug treatments of OCI-AML3 cells. **D.** Mean total base pairs covered by p53 ChIP-seq peaks (peaks present in at least 2 out of 3 replicates; mean of 2-3 replicates) after the indicated drug treatments of OCI-AML3 cells. **E.** Heat map of p53 ChIP-seq peaks (peaks present in at least 2 out of 3 replicates) at 252 synergy up genes (rows) after the indicated drug treatments (columns) of OCI-AML3 cells.

**Supp Figure 5.**
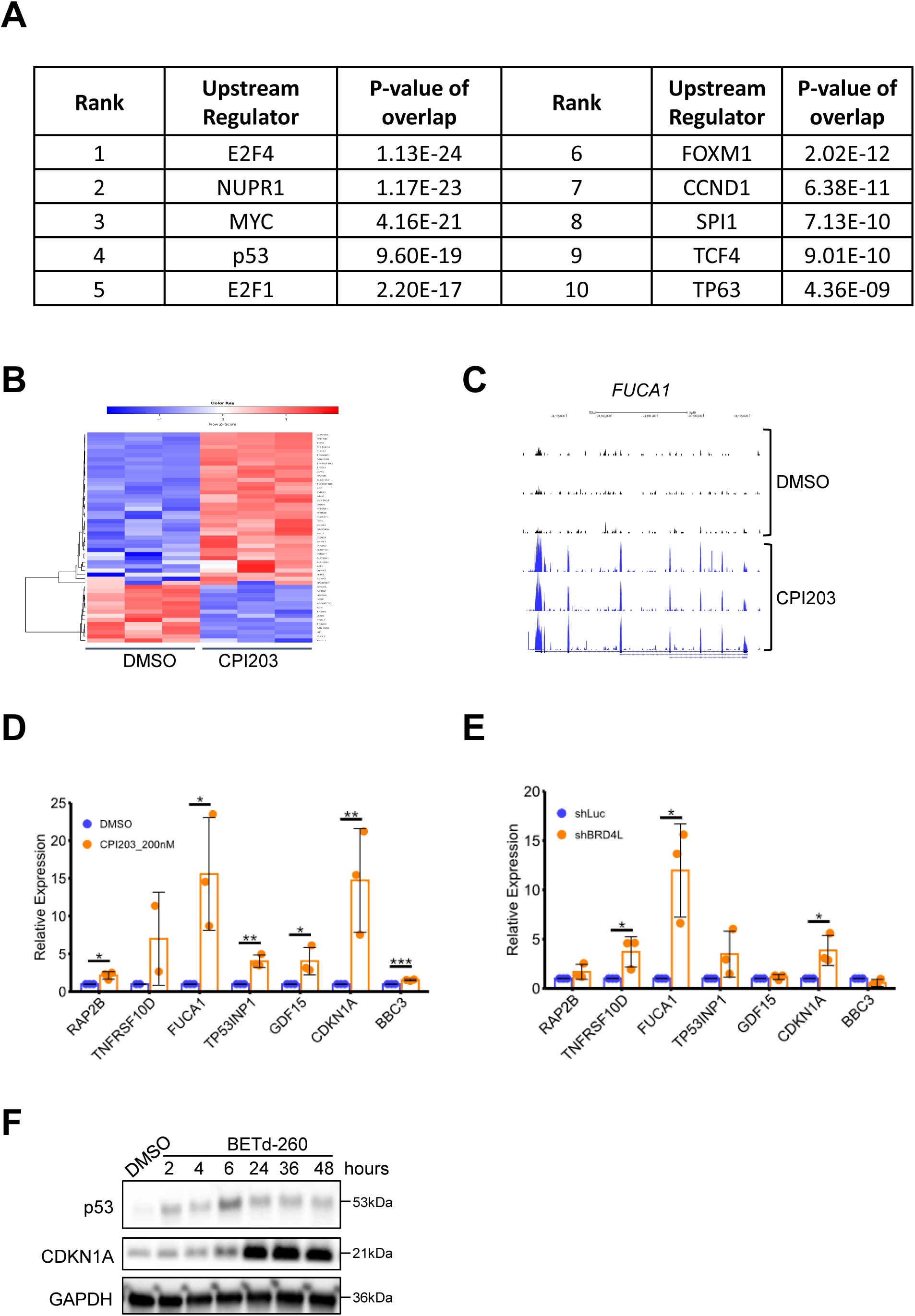
Inhibition of BET family proteins activates p53. **A.** Ingenuity pathway analysis (IPA) identifies p53 as a candidate regulator of genes differentially expressed between control and CPI203-treated AML3 cells. p53 is predicted to be activated, with an activation z-score 5.802 (fisher’s exact test). **B.** Heat map of all significant changes in gene expression of known p53 target genes (from (Fischer, 2017)) in RNA-seq data of OCI-AML3 cells, control (DMSO) vs CPI203 treated AML3 cells (3 replicates of each). **C.** RNA-seq sequence tags from control (DMSO) and CPI203-treated AML3 cells aligned to *FUCA1* gene (3 replicates of each). **D.** qPCR analysis of indicated p53 target genes in samples from Figure 3E (***=p≤0.001, **=p≤0.01, *=p≤0.05, two tailed unpaired t-test, Means +/− SD are shown, n=3). **E.** qPCR analysis of indicated p53 target genes in OCI-AML3 cells expressing small hairpin RNA(shRNA) specific for luciferase gene (shLuc, control) or BRD4L (shBRD4L) (*=p≤0.05, two tailed unpaired t-test, Means +/− SD are shown, n=3). **F.** Western blot of p53 and CDKN1A in OCI-AML3 cells treated with BETd-260 at 5nM for different times.

**Supp Figure 6.**
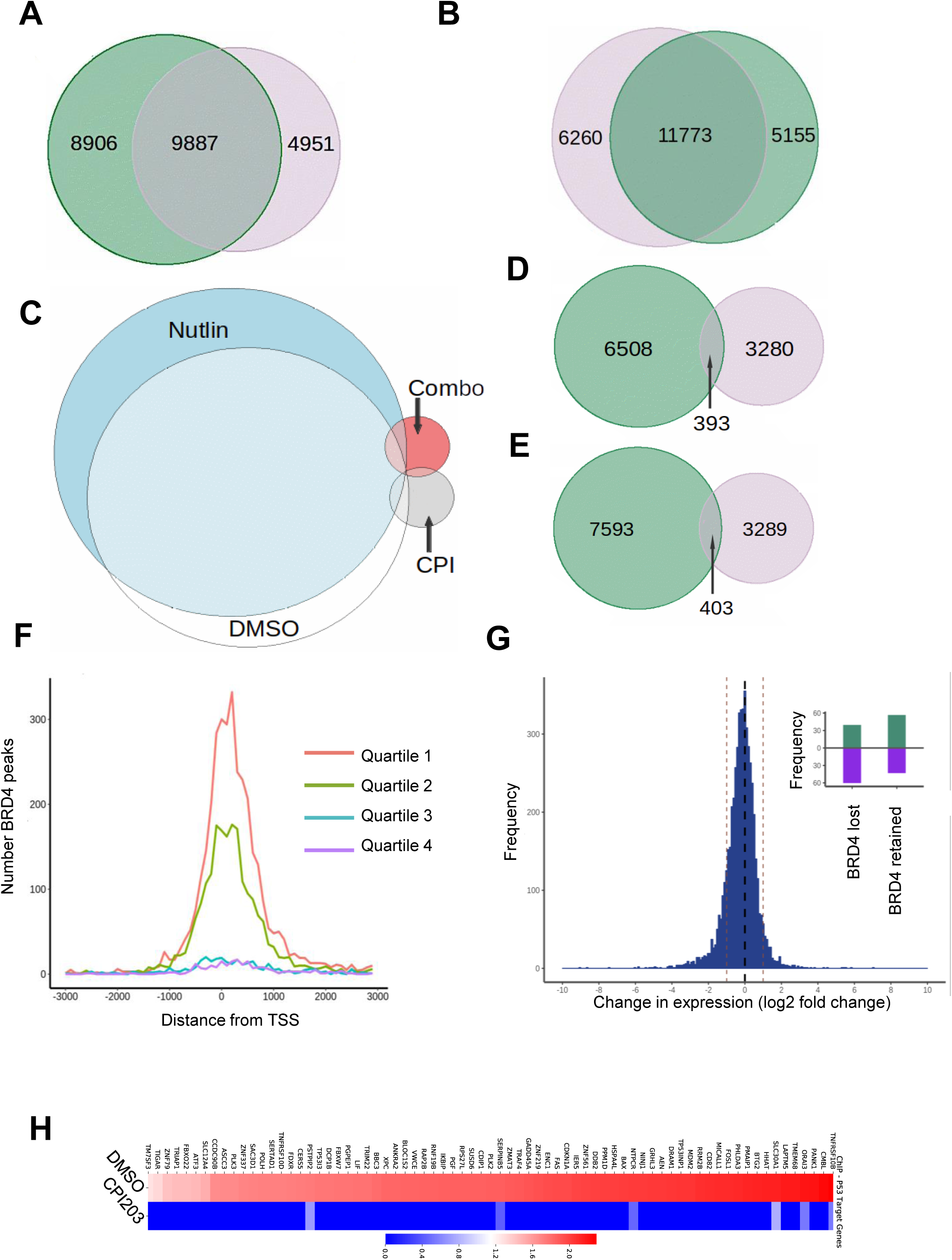
BRD4 ChIP-seq data. **A.** Venn diagram showing overlap of 2 replicates of BRD4 ChIP-seq peaks in OCI-AML3 cells treated with DMSO **B.** Venn diagram showing overlap of 2 replicates of BRD4 ChIP-seq peaks in OCI-AML3 cells treated with nutlin-3 **C.** Venn diagram showing overlap of BRD4 ChIP-seq peaks (peaks present in 2 out of 2 replicates) in OCI-AML3 cells treated with DMSO, nutlin-3, CPI203 or combo. **D.** Venn diagram showing overlap of 2 replicates of BRD4 ChIP-seq peaks in OCI-AML3 cells treated with CPI203 **E.** Venn diagram showing overlap of 2 replicates of BRD4 ChIP-seq peaks in OCI-AML3 cells treated with CPI203 and nutlin-3. **F.** Histogram of number of BRD4 ChIP-seqs centered around gene TSS (0) for genes in the highest (quartile 1) to lowest (quartile 4) quartiles of expression in DMSO treated OCI-AML3 cells. **G.** The histogram shows the spread of change in expression for genes losing BRD4 binding after treatment with CPI203. Loss of BRD4 on CPI203 treatment was associated with a decrease in gene expression (p<0.05). Inset figure: BRD4 peaks overlapping TSS were divided into two groups, those losing BRD4 on CPI203 treatment and those retaining BRD4. Plotted is the % change in expression after CPI203 treatment for the genes within each group. **H.** Heat map of BRD4 ChIP-seq peaks in known p53 target genes from Control (DMSO) and CPI203 treatment. Values represent the aggregated read counts associated with each gene across samples in log scale.

**Supp Figure 7.**
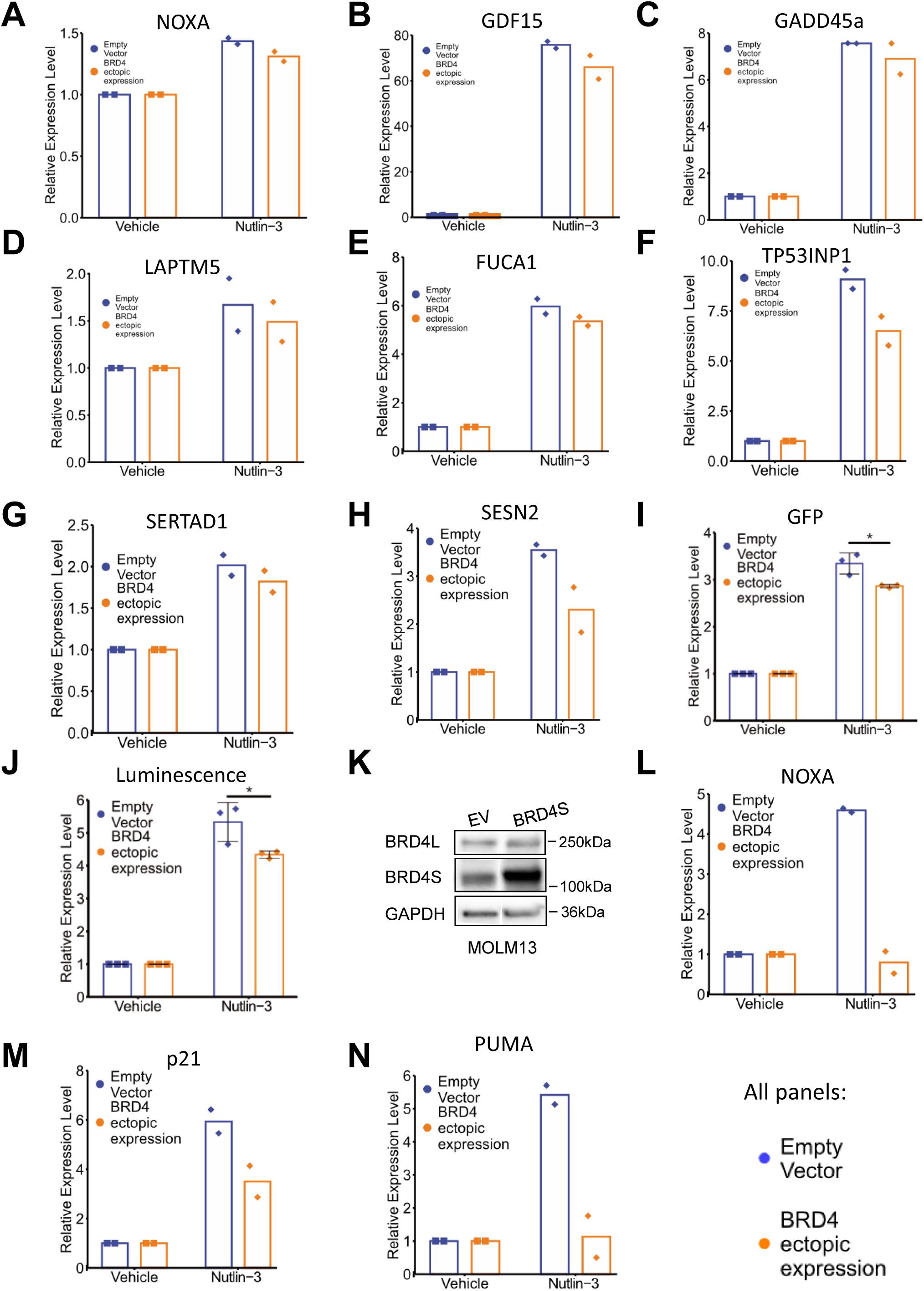
BRD4 represses p53 target genes. **A.** qPCR analysis of *NOXA* in OCI-AML3 cells ectopically expressing BRD4S, in absence or presence of nutlin-3 (2 replicates of each). **B.** qPCR analysis of *GDF15* in OCI-AML3 cells ectopically expressing BRD4S, in absence or presence of nutlin-3 (2 replicates of each). **C.** qPCR analysis of *GADD45a* in OCI-AML3 cells ectopically expressing BRD4S, in absence or presence of nutlin-3 (2 replicates of each). **D.** qPCR analysis of *LAPTM5* in OCI-AML3 cells ectopically expressing BRD4S, in absence or presence of nutlin-3 (2 replicates of each). **E.** qPCR analysis of *FUCA1* in OCI-AML3 cells ectopically expressing BRD4S, in absence or presence of nutlin-3 (2 replicates of each). **F.** qPCR analysis of *TP53INP1* in OCI-AML3 cells ectopically expressing BRD4S, in absence or presence of nutlin-3 (2 replicates of each). **G.** qPCR analysis of *SERTAD1* in OCI-AML3 cells ectopically expressing BRD4S, in absence or presence of nutlin-3 (2 replicates of each). **H.** qPCR analysis of *SESN2* in OCI-AML3 cells ectopically expressing BRD4S, in absence or presence of nutlin-3 (2 replicates of each). **I.** GFP measurement in OCI-AML3 cells ectopically expressing BRD4S, in absence or presence of nutlin-3(*=p≤0.05 by a two-tailed unpaired t-test, n=3). **J.** Luminescence measurement in OCI-AML3 cells ectopically expressing BRD4S, in absence or presence of nutlin-3(*=p≤0.05 by a two-tailed unpaired t-test, n=3). **K.** Western blot for BRD4 in MOLM13 cells ectopically expressing the short isoform of BRD4 (BRD4S). BRD4L is the long isoform. **L.** qPCR analysis of *NOXA* in OCI-AML3 cells ectopically expressing BRD4S, in absence or presence of nutlin-3 (**=p≤0.01) (2 replicates of each). **M.** qPCR analysis of *CDKN1A* in OCI-AML3 cells ectopically expressing BRD4S, in absence or presence of nutlin-3 (2 replicates of each). **N.** qPCR analysis of *PUMA* in OCI-AML3 cells ectopically expressing BRD4S, in absence or presence of nutlin-3 (*=p≤0.05) (2 replicates of each).

## References

Abraham, S.A., Hopcroft, L.E., Carrick, E., Drotar, M.E., Dunn, K., Williamson, A.J., Korfi, K., Baquero, P., Park, L.E., Scott, M.T., et al. (2016). Dual targeting of p53 and c-MYC selectively eliminates leukaemic stem cells. Nature 534, 341–346.

Albrecht, B.K., Gehling, V.S., Hewitt, M.C., Vaswani, R.G., Cote, A., Leblanc, Y., Nasveschuk, C.G., Bellon, S., Bergeron, L., Campbell, R., et al. (2016). Identification of a Benzoisoxazoloazepine Inhibitor (CPI-0610) of the Bromodomain and Extra-Terminal (BET) Family as a Candidate for Human Clinical Trials. J Med Chem 59, 1330–1339.

Alqahtani, A., Choucair, K., Ashraf, M., Hammouda, D.M., Alloghbi, A., Khan, T., Senzer, N., and Nemunaitis, J. (2019). Bromodomain and extra-terminal motif inhibitors: a review of preclinical and clinical advances in cancer therapy. Future Sci OA 5, FSO372.

Amorim, S., Stathis, A., Gleeson, M., Iyengar, S., Magarotto, V., Leleu, X., Morschhauser, F., Karlin, L., Broussais, F., Rezai, K., et al. (2016). Bromodomain inhibitor OTX015 in patients with lymphoma or multiple myeloma: a dose-escalation, open-label, pharmacokinetic, phase 1 study. Lancet Haematol 3, e196–204.

Andreeff, M., Kelly, K.R., Yee, K., Assouline, S., Strair, R., Popplewell, L., Bowen, D., Martinelli, G., Drummond, M.W., Vyas, P., et al. (2016). Results of the Phase I Trial of RG7112, a Small-Molecule MDM2 Antagonist in Leukemia. Clin Cancer Res 22, 868–876.

Bankar, A., and Gupta, V. (2020). Investigational non-JAK inhibitors for chronic phase myelofibrosis. Expert Opin Investig Drugs 29, 461–474.

Bansal, H., Kornblau, S., Yihua, Q., Coombes, K., Panneerdoss, S., Karnad, A., Weitman, S., Tomlinson, G., Bansal, S., and Iyer, S. (2017). Overexpression of BRD4 Is an Adverse Prognostic Factor in Acute Myeloid Leukemia. In American Society of Hematology Annual Meeting 2017 (Blood), pp. 3794.

Berthon, C., Raffoux, E., Thomas, X., Vey, N., Gomez-Roca, C., Yee, K., Taussig, D.C., Rezai, K., Roumier, C., Herait, P., et al. (2016). Bromodomain inhibitor OTX015 in patients with acute leukaemia: a dose-escalation, phase 1 study. Lancet Haematol 3, e186–195.

Chaidos, A., Caputo, V., and Karadimitris, A. (2015). Inhibition of bromodomain and extra-terminal proteins (BET) as a potential therapeutic approach in haematological malignancies: emerging preclinical and clinical evidence. Ther Adv Hematol 6, 128–141.

Chan, I.T., and Gilliland, D.G. (2004). Oncogenic K-ras in mouse models of myeloproliferative disease and acute myeloid leukemia. Cell Cycle 3, 536–537.

Chou, T.C., and Talalay, P. (1984). Quantitative analysis of dose-effect relationships: the combined effects of multiple drugs or enzyme inhibitors. Adv Enzyme Regul 22, 27–55.

Conrad, R.J., Fozouni, P., Thomas, S., Sy, H., Zhang, Q., Zhou, M.M., and Ott, M. (2017). The Short Isoform of BRD4 Promotes HIV-1 Latency by Engaging Repressive SWI/SNF Chromatin-Remodeling Complexes. Mol Cell 67, 1001–1012 e1006.

Cuella-Martin, R., Oliveira, C., Lockstone, H.E., Snellenberg, S., Grolmusova, N., and Chapman, J.R. (2016). 53BP1 Integrates DNA Repair and p53-Dependent Cell Fate Decisions via Distinct Mechanisms. Mol Cell 64, 51–64.

Dawson, M.A., Prinjha, R.K., Dittmann, A., Giotopoulos, G., Bantscheff, M., Chan, W.I., Robson, S.C., Chung, C.W., Hopf, C., Savitski, M.M., et al. (2011). Inhibition of BET recruitment to chromatin as an effective treatment for MLL-fusion leukaemia. Nature 478, 529–533.

Delgado, M.D., and Leon, J. (2010). Myc roles in hematopoiesis and leukemia. Genes Cancer 1, 605–616.

Ding, Q., Zhang, Z., Liu, J.J., Jiang, N., Zhang, J., Ross, T.M., Chu, X.J., Bartkovitz, D., Podlaski, F., Janson, C., et al. (2013). Discovery of RG7388, a potent and selective p53-MDM2 inhibitor in clinical development. J Med Chem 56, 5979–5983.

Dombret, H., and Gardin, C. (2016). An update of current treatments for adult acute myeloid leukemia. Blood 127, 53–61.

Filippakopoulos, P., Qi, J., Picaud, S., Shen, Y., Smith, W.B., Fedorov, O., Morse, E.M., Keates, T., Hickman, T.T., Felletar, I., et al. (2010). Selective inhibition of BET bromodomains. Nature 468, 1067–1073.

Fischer, M. (2017). Census and evaluation of p53 target genes. Oncogene 36, 3943–3956.

Fuchs, S.Y., Adler, V., Buschmann, T., Yin, Z., Wu, X., Jones, S.N., and Ronai, Z. (1998). JNK targets p53 ubiquitination and degradation in nonstressed cells. Genes Dev 12, 2658–2663.

Ianevski, A., Giri, A.K., and Aittokallio, T. (2020). SynergyFinder 2.0: visual analytics of multi-drug combination synergies. Nucleic Acids Res.

Jang, M.K., Mochizuki, K., Zhou, M., Jeong, H.S., Brady, J.N., and Ozato, K. (2005). The bromodomain protein Brd4 is a positive regulatory component of P-TEFb and stimulates RNA polymerase II-dependent transcription. Mol Cell 19, 523–534.

Kastenhuber, E.R., and Lowe, S.W. (2017). Putting p53 in Context. Cell 170, 1062–1078.

Keeshan, K., He, Y., Wouters, B.J., Shestova, O., Xu, L., Sai, H., Rodriguez, C.G., Maillard, I., Tobias, J.W., Valk, P., et al. (2006). Tribbles homolog 2 inactivates C/EBPalpha and causes acute myelogenous leukemia. Cancer Cell 10, 401–411.

Khoo, K.H., Verma, C.S., and Lane, D.P. (2014). Drugging the p53 pathway: understanding the route to clinical efficacy. Nat Rev Drug Discov 13, 217–236.

Kirschner, K., Samarajiwa, S.A., Cairns, J.M., Menon, S., Perez-Mancera, P.A., Tomimatsu, K., Bermejo-Rodriguez, C., Ito, Y., Chandra, T., Narita, M., et al. (2015). Phenotype specific analyses reveal distinct regulatory mechanism for chronically activated p53. PLoS Genet 11, e1005053.

Klco, J.M., Spencer, D.H., Lamprecht, T.L., Sarkaria, S.M., Wylie, T., Magrini, V., Hundal, J., Walker, J., Varghese, N., Erdmann-Gilmore, P., et al. (2013). Genomic and epigenomic landscapes of adult de novo acute myeloid leukemia. N Engl J Med 368, 2059–2074.

Lu, J., Qian, Y., Altieri, M., Dong, H., Wang, J., Raina, K., Hines, J., Winkler, J.D., Crew, A.P., Coleman, K., et al. (2015). Hijacking the E3 Ubiquitin Ligase Cereblon to Efficiently Target BRD4. Chem Biol 22, 755–763.

Maganti, H.B., Jrade, H., Cafariello, C., Manias Rothberg, J.L., Porter, C.J., Yockell-Lelievre, J., Battaion, H.L., Khan, S.T., Howard, J.P., Li, Y., et al. (2018). Targeting the MTF2-MDM2 Axis Sensitizes Refractory Acute Myeloid Leukemia to Chemotherapy. Cancer Discov 8, 1376–1389.

Matsuo, Y., MacLeod, R.A., Uphoff, C.C., Drexler, H.G., Nishizaki, C., Katayama, Y., Kimura, G., Fujii, N., Omoto, E., Harada, M., et al. (1997). Two acute monocytic leukemia (AML-M5a) cell lines (MOLM-13 and MOLM-14) with interclonal phenotypic heterogeneity showing MLL-AF9 fusion resulting from an occult chromosome insertion, ins(11;9)(q23;p22p23). Leukemia 11, 1469–1477.

Mertz, J.A., Conery, A.R., Bryant, B.M., Sandy, P., Balasubramanian, S., Mele, D.A., Bergeron, L., and Sims, R.J., 3rd (2011). Targeting MYC dependence in cancer by inhibiting BET bromodomains. Proc Natl Acad Sci U S A 108, 16669–16674.

Minzel, W., Venkatachalam, A., Fink, A., Hung, E., Brachya, G., Burstain, I., Shaham, M., Rivlin, A., Omer, I., Zinger, A., et al. (2018). Small Molecules Co-targeting CKIalpha and the Transcriptional Kinases CDK7/9 Control AML in Preclinical Models. Cell 175, 171–185 e125.

Moros, A., Rodriguez, V., Saborit-Villarroya, I., Montraveta, A., Balsas, P., Sandy, P., Martinez, A., Wiestner, A., Normant, E., Campo, E., et al. (2014). Synergistic antitumor activity of lenalidomide with the BET bromodomain inhibitor CPI203 in bortezomib-resistant mantle cell lymphoma. Leukemia 28, 2049–2059.

Pan, R., Ruvolo, V., Mu, H., Leverson, J.D., Nichols, G., Reed, J.C., Konopleva, M., and Andreeff, M. (2017). Synthetic Lethality of Combined Bcl-2 Inhibition and p53 Activation in AML: Mechanisms and Superior Antileukemic Efficacy. Cancer Cell 32, 748–760 e746.

Papaemmanuil, E., Gerstung, M., Bullinger, L., Gaidzik, V.I., Paschka, P., Roberts, N.D., Potter, N.E., Heuser, M., Thol, F., Bolli, N., et al. (2016). Genomic Classification and Prognosis in Acute Myeloid Leukemia. N Engl J Med 374, 2209–2221.

Prokocimer, M., Molchadsky, A., and Rotter, V. (2017). Dysfunctional diversity of p53 proteins in adult acute myeloid leukemia: projections on diagnostic workup and therapy. Blood 130, 699–712.

Roe, J.S., and Vakoc, C.R. (2016). The Essential Transcriptional Function of BRD4 in Acute Myeloid Leukemia. Cold Spring Harb Symp Quant Biol 81, 61–66.

Sakamaki, J.I., Wilkinson, S., Hahn, M., Tasdemir, N., O’Prey, J., Clark, W., Hedley, A., Nixon, C., Long, J.S., New, M., et al. (2017). Bromodomain Protein BRD4 Is a Transcriptional Repressor of Autophagy and Lysosomal Function. Mol Cell 66, 517–532 e519.

Schmidt, D., Wilson, M.D., Spyrou, C., Brown, G.D., Hadfield, J., and Odom, D.T. (2009). ChIP-seq: using high-throughput sequencing to discover protein-DNA interactions. Methods 48, 240–248.

Somervaille, T.C., and Cleary, M.L. (2006). Identification and characterization of leukemia stem cells in murine MLL-AF9 acute myeloid leukemia. Cancer Cell 10, 257–268.

Stewart, H.J., Horne, G.A., Bastow, S., and Chevassut, T.J. (2013). BRD4 associates with p53 in DNMT3A-mutated leukemia cells and is implicated in apoptosis by the bromodomain inhibitor JQ1. Cancer Med 2, 826–835.

Vassilev, L.T., Vu, B.T., Graves, B., Carvajal, D., Podlaski, F., Filipovic, Z., Kong, N., Kammlott, U., Lukacs, C., Klein, C., et al. (2004). In vivo activation of the p53 pathway by small-molecule antagonists of MDM2. Science 303, 844–848.

Watts, J., and Nimer, S. (2018). Recent advances in the understanding and treatment of acute myeloid leukemia. F1000Res 7.

Wu, S.Y., Lee, A.Y., Hou, S.Y., Kemper, J.K., Erdjument-Bromage, H., Tempst, P., and Chiang, C.M. (2006). Brd4 links chromatin targeting to HPV transcriptional silencing. Genes Dev 20, 2383–2396.

Wu, S.Y., Lee, A.Y., Lai, H.T., Zhang, H., and Chiang, C.M. (2013). Phospho switch triggers Brd4 chromatin binding and activator recruitment for gene-specific targeting. Mol Cell 49, 843–857.

Xu, Z., Sharp, P.P., Yao, Y., Segal, D., Ang, C.H., Khaw, S.L., Aubrey, B.J., Gong, J., Kelly, G.L., Herold, M.J., et al. (2016). BET inhibition represses miR17-92 to drive BIM-initiated apoptosis of normal and transformed hematopoietic cells. Leukemia 30, 1531–1541.

Zhang, J., Kong, G., Rajagopalan, A., Lu, L., Song, J., Hussaini, M., Zhang, X., Ranheim, E.A., Liu, Y., Wang, J., et al. (2017). p53-/- synergizes with enhanced NrasG12D signaling to transform megakaryocyte-erythroid progenitors in acute myeloid leukemia. Blood 129, 358–370.

Zhao, Z., Zuber, J., Diaz-Flores, E., Lintault, L., Kogan, S.C., Shannon, K., and Lowe, S.W. (2010). p53 loss promotes acute myeloid leukemia by enabling aberrant self-renewal. Genes Dev 24, 1389–1402.

Zhou, B., Hu, J., Xu, F., Chen, Z., Bai, L., Fernandez-Salas, E., Lin, M., Liu, L., Yang, C.Y., Zhao, Y., et al. (2018). Discovery of a Small-Molecule Degrader of Bromodomain and Extra-Terminal (BET) Proteins with Picomolar Cellular Potencies and Capable of Achieving Tumor Regression. J Med Chem 61, 462–481.

Zuber, J., Shi, J., Wang, E., Rappaport, A.R., Herrmann, H., Sison, E.A., Magoon, D., Qi, J., Blatt, K., Wunderlich, M., et al. (2011). RNAi screen identifies Brd4 as a therapeutic target in acute myeloid leukaemia. Nature 478, 524–528.

